# Novel Role of Endothelial CD45 in Regulating Endothelial-to-Mesenchymal Transition in Atherosclerosis

**DOI:** 10.1101/2024.09.03.610974

**Authors:** Qianman Peng, Kulandaisamy Arulsamy, Yao Wei Lu, Bo Zhu, Bandana Singh, Kui Cui, Jill Wylie-Sears, Marina V. Malovichko, Scott Wong, Douglas B. Cowan, Masanori Aikawa, Sanjay Srivastava, Da-Zhi Wang, Joyce Bischoff, Kaifu Chen, Hong Chen

## Abstract

**Background:** The protein tyrosine phosphatase CD45 is expressed in all nucleated cells of the hematopoietic system and in mitral valve endothelial cells (ECs) undergoing endothelial-to-mesenchymal transition (EndoMT). Our recent work indicated that activation of endogenous CD45 in human endothelial colony-forming cells (ECFCs) induced expression of multiple EndoMT marker genes. We hypothesized that CD45 may contribute to atherosclerosis; however, detailed molecular mechanisms underlying how CD45 may contribute to EndoMT and the impact of therapeutic manipulation of CD45 expression in atherosclerosis are unknown.

**Methods:** We generated a tamoxifen-inducible EC-specific CD45-deficient mouse strain (EC-iCD45KO) on an ApoE-deficient (WT/ApoE^−/−^) background and fed them a Western diet (WD) to produce atherosclerosis. We enriched mouse aortic ECs with anti-CD31 beads to perform single-cell RNA sequencing. Cellular, biochemical and molecular approaches were used to investigate the effect of endothelial CD45-specific deletion on EndoMT and lesion development in an ApoE^−/−^ mouse model of atherosclerosis.

**Results:** EC-iCD45KO mice showed reductions in lesion development, plaque macrophage infiltration, and expression of cell adhesion molecules when compared to WT/ApoE^−/−^ controls. Single-cell RNA sequencing revealed that loss of endothelial CD45 decreases EndoMT marker expression and TGF-β signaling in atherosclerotic mice, which is associated with reduction of lesions. Mechanistically, CD45 loss increases Fibroblast Growth Factor Receptor 2 (FGFR2) expression in mouse aortic ECs and Krüppel-like Factor 2 (KLF2) expression in the aortic root. Endothelial CD45-deficiency also inhibits EndoMT and TGFβ signaling in atherosclerosis.

**Conclusions:** Our findings demonstrate that genetic depletion of endothelial CD45 protects against EndoMT-driven atherosclerosis by promoting FGFR2 and KLF2 expression while inhibiting Transforming Growth Factor beta (TGFβ) signaling and EndoMT. Consequently, targeting endothelial CD45 may represent a novel therapeutic strategy to reduce EndoMT in atherosclerosis.

## INTRODUCTION

Accumulating evidence indicates that CD45 (*PTPRC*), the prototypic receptor-like protein tyrosine phosphatase (PTP), is an essential regulator of signal transduction in hematopoietic cells. It dephosphorylates phosphotyrosine residues to dampen cell signaling^1^. While CD45 is not commonly expressed in endothelial cells (ECs), lung ECs have been found to express CD31 and CD45 in a pulmonary arterial hypertension rat model^2^. Notably, we recently reported that activation of the endogenous CD45 promoter in human endothelial colony forming cells (ECFCs) induced expression of endothelial-to-mesenchymal transition (EndoMT) markers and increased cell migration, contraction of collagen gels, and permeability consistent with EndoMT^3^. In addition, a CD45 phosphatase inhibitor blocked EndoMT in Transforming Growth Factor beta 1 (TGFβ1)-treated ovine mitral valve ECs^4^. These findings led us to speculate that endothelial CD45-specific deletion may prevent EndoMT-driven atherosclerosis.

Atherosclerosis is the leading cause of morbidity and mortality in the United States and most developed countries^5^. Endothelial dysfunction caused by cholesterol and oxidized phospholipids in lipoproteins (*e.g.*, oxidized Low Density Lipoprotein or oxLDL) are major atherosclerosis-initiating events^6^. Trapped lipoproteins activate ECs and vascular smooth muscle cells (VSMCs) to increase expression of adhesion molecules and chemokines^7^. This leads to recruitment of monocytes/lymphocytes into the arterial wall^8^. Lesion progression involves the migration and proliferation of resident smooth muscle cells (SMCs)^9^, extracellular matrix deposition^8^, and lipid or necrotic core formation^10^. In advanced stages of atherosclerosis, there is increased endothelial dysfunction and advanced plaque ruptures associated with thin, collagen-poor fibrous caps^11^ that can result in myocardial infarctions and ischemic strokes.

EndoMT is an incremental process where ECs progressively lose endothelial-specific markers and properties while gaining mesenchymal attributes^12^. ECs can be distinguished by expression of specific endothelial genes in normal physiological conditions. For example, ECs express platelet/EC adhesion molecule-1 (CD31/PECAM-1), vascular endothelial (VE)-cadherin, and von Willebrand factor, which are reduced during EndoMT^13^. At the same time, transitioning ECs express mesenchymal markers such as αSMA, collagen, and SM22^14^. While EndoMT plays a fundamental role in forming cardiac valves of the heart development, there is mounting evidence that EndoMT is involved in atherosclerosis in adults^15,16^.

TGFβ signaling is a central player in driving EndoMT during atheroma progression^17^ but the processes leading to its activation are poorly defined. TGFβ signaling in ECs during EndoMT results in expression of pro-inflammatory chemokines and cytokines as well as their receptors^17^, leukocyte adhesion molecules (*e.g.*, ICAM-1 and VCAM-1)^18^, and extracellular matrix (ECM) components linked to inflammation^8^. In contrast, a non-inflammatory EC phenotype can be preserved through FGFR signaling, which counteracts TGFβ-driven EndoMT^19^. Reductions in endothelial FGFR-mediated signaling enhances TGFβ signaling leading to EndoMT^20^. Endothelial FGFR1 and the regulatory transcription factor Krüppel-like Factor 2 (KLF2) are reduced by inflammation^21^, whereas FGFR2 activation can enhance autocrine FGF2 production, which has been shown to improve cardiac function in infarcted mice^22^.

The objective of this study was to establish if endothelial CD45-specific deletion could prevent EndoMT in atherosclerosis. To do this, we generated tamoxifen-inducible EC-specific CD45-deficient mice (EC-iCD45KO) on an ApoE^−/−^background then fed these mice an atherosclerotic Western diet (WD). We performed single-cell RNA sequencing (scRNA-seq) and biochemical experiments to assess endothelial CD45-mediated TGFβ- and FGF-receptor signaling *in vitro* and *in vivo*. Our findings demonstrate that loss of endothelial CD45 protects against EndoMT-driven atherosclerosis, while simultaneously promoting FGFR2 and KLF2 expression and inhibiting TGFβ signaling and EndoMT marker expression. These studies strongly indicate that endothelial CD45 may represent a novel drug target for the treatment of atherosclerosis.

## MATERIALS AND METHODS

Additional Materials and Methods are available in the Supplemental Materials and Methods and Supplemental Tables.

### Genetic Mouse Models and Aortic Lesion Evaluation

All animal procedures were performed in compliance with institutional guidelines, and mouse protocols were approved by the Institutional Animal Care and Use Committee (IACUC) of Boston Children’s Hospital. Both male and female mice were used, including C57BL/6 mice (Jax stock #00664), ApoE^−/−^ mice (Jax stock #002052), and EC-specific Cre deleter mice (iCDH5CreERT2) (The Jackson Laboratory). Please see the Supplemental Materials and Methods for a description of the breeding scheme. Tamoxifen-inducible deletion of endothelial CD45 on an ApoE^−/−^ background (EC-iCD45KO/ApoE^−/−^) was accomplished by performing 7 injections every other day as previously described^23^. For controls, C57BL/6 iCDH5 CreERT2 mice were denoted as wild-type (WT). These mice were crossed onto an ApoE^−/−^ background to obtain control ApoE^−/−^/WT (WT/ApoE^−/−^) mice. WT/ApoE^−/−^ and EC-iCD45KO/ApoE^−/−^ were fed a Western diet (WD) for 12-16 weeks beginning at 8 months of age. We evaluated atherosclerotic lesions at the aortic sinus, abdominal aorta, and thoracic aorta. Mice were perfused, dissected, and subjected to quantification of atherosclerosis as previously described^24^.

### Cell Culture

To isolate murine aortic ECs (MAECs), aortas were collected and washed twice with PBS at 4°C and then stripped of fat and connective tissue^23,24^. Aortas were cut into 3 mm long sections, and segments were put on the Matrigel-coated (Corning) plate with EC medium^25^. After 4 days, vascular networks were visible under the light microscope and tissue segments were removed. ECs were detached and cultured in fresh endothelial medium. The identity of isolated ECs was confirmed by immunofluorescence staining for the EC marker CD31. A full list of reagents including antibodies and primers is included in the Supplemental Tables. Cultured MAECs isolated from EC-iCD45KO/ApoE^−/−^ iCDH5-Cre mice were treated with 5 µM tamoxifen for 3 days to induce deletion of CD45. MAECs isolated from ApoE^−/−^ mice were treated in parallel with tamoxifen to serve as controls. Cells were treated with 100 µg/mL oxLDL or 10 ng/mL TGFβ1 while maintaining 2 µM tamoxifen in the medium.

### Aortic Sample Preparation for scRNA-seq

Aortic cells were isolated as described previously^23^. Mouse aortas were cut into fine pieces and placed in 5 mL of enzyme solution at 37°C. Supernatants were collected every 10 min until all pieces were digested (Enzyme solution: 5 mL of DMEM, Collagenase type I: 5mg/mL (Gibco), Collagenase type IV: 5mg/mL (Gibco), Liberase:5mg/mL (Roche)). Supernatants were filtered with a 40-µm strainer, centrifuged at 400g for 5 mins, and the cell pellet was suspended in 90 µL of buffer containing 0.5% BSA and 2 mmol/L EDTA in PBS. Then, 20 µL of CD31 microbeads (Miltenyi Biotec) was added to the suspension at 4° C for 15 mins. Microbeads were passed through an MS column (Miltenyi Biotec). Cells were processed for scRNA-seq with the 10x Genomics platform and sequencing was performed by MedGenome (**Figure S1**).

### Profiling Single-Cell Transcriptomes Using 10X scRNA-sequencing

The raw sequence reads from WT/ApoE^−/−^ and EC-iCD45KO/ApoE^−/−^ samples were obtained using 10X Genomics scRNA-seq and processed independently. These reads were aligned against the mouse reference genome (mm10) using Cell Ranger^26^ (version 7.1.0). This pipeline generated a gene expression count matrix, with rows representing genes and columns representing individual cells. Subsequently, we employed the Seurat R package (version 5.1.0)^27^ for downstream analysis. Low-quality cells were excluded if they had fewer than 500 or more than 4500 expressed genes, or if the percentage of mitochondrial or ribosomal reads exceeded 5% or 16%, respectively. For each sample, doublet cells identified by the DoubletFinder^28^ program were removed. The remaining high-quality cells were then subjected to log normalization, data scaling, principal component analysis (PCA), and cell clustering. Cell clustering and sub-clustering were performed using the FindClusters and FindSubCluster functions, respectively. Marker genes for each cluster were identified using the FindAllMarkers function, with a minimum percentage of expressed cells set at 25% and a minimum log2 fold change of 0.25. These marker genes were used to annotate cell types based on known markers from public databases^29,30^ and the literature^23^. Differentially expressed genes between WT/ApoE^−/−^ and EC-iCD45KO/ApoE^−/−^ samples were analyzed by grouping cells from each sample by the interested cell types (EndoMT1 and EndoMT2) and running the FindMarkers function. For pathway enrichment analysis, we utilized the clusterProfiler R package^31^ to explore the Gene Ontology (GO) pathways of differentially regulated genes. In case of data visualization, we utilized the DimPlot for UMAP plots, and VlnPlot and DotPlot functions for plotting gene expression levels across different cell types between samples. The Wilcoxon non-parametric test was used to calculate p-values for single-cell gene expression analysis, and a proportion test was applied to assess the statistical significance of differences in cell type proportions between WT/ApoE^−/−^ and EC-iCD45KO/ApoE^−/−^ groups. We utilized the SCIG tool^32^ to characterize cell identity genes across cell types in both groups, followed by Gene Ontology (GO) pathway analysis to investigate their functional roles. Further, we analyzed human single-cell RNA-seq data from healthy controls and atherosclerosis samples^33,34^ to assess the expression of CD45/PTPRC and other endothelial functional genes such as CDH5, EMCN, and FLT1 in Pecam1-expressing cells.

### Single-Cell Trajectory, RNA Velocity, and Cell–Cell Communication Analyses

To investigate the role of CD45 in the EndoMT process, we performed single-cell cellular dynamics analysis using RNA velocity, implemented with the scVelo Python package. In this approach, the future states of individual cells were modeled based on the ratio of unspliced and spliced mRNA transcripts abundance^35^. Trajectory inference analysis for each sample was conducted using the R package monocle3^36^. Changes in gene expression along trajectories were visualized with the R ComplexHeatmap^37^ package. We investigated cell-cell communication by identifying potential ligands in sender cells and their corresponding target genes in receiver cells using the nichenetr package^38^. To understand which cell-cell communication signals regulate the EndoMT process, we specifically performed the analysis by considering EndoMT cells as sender cells and ECs as receiver cells.

### Hematoxylin and Eosin Staining

Cryostat sections of the mouse aortic root, abdominal aorta, and brachiocephalic artery (BCA) were washed in PBS for 5 mins, then fixed in 4% paraformaldehyde for 15 mins. Slides were stained with hematoxylin for 3 mins, followed by running tap water washes for 10 mins. Slices were then dipped in eosin working solution for 30 seconds, rinsed with tap water, and dehydrated using graded ethanol (95% and 100% ethanol), followed by treatment with xylenes. Slices were mounted in synthetic resin as previously described^39^.

### Flow Cytometry

Flow cytometry analysis of aortic cells from normal diet (ND) ApoE^−/−^ and WD diet ApoE^−/−^(16-20 weeks) was performed on cells isolated as described above. Cell pellets were resuspended in FACS buffer (2% HBSS, 0.004% FBS, 2mM EDTA in PBS) at 4°C for 30 min to block nonspecific binding of antibodies to Fc receptors. Cells were incubated with antibodies for 30 mins and flow cytometry analyses were performed using a BD instrument.

### Oil Red O Staining

Cryostat sections were fixed in 4% paraformaldehyde^23,39^. Slides were washed with PBS, and rinsed with 60% isopropanol, followed by staining with freshly prepared 0.5% Oil Red O solution in isopropanol for 20 mins at 37°C^39^. Slices were then put in 60% isopropanol for 30 seconds, followed by 3 washes in water, and dried slides were mounted with glycerin. Imaging was performed using a Zeiss LSM880 microscope and analyzed with ZEN-Lite software and NIH ImageJ software.

Whole aortae were dissected symmetrically, pinned to parafilm for *en face* exposure and fixed in formalin for 24 hrs. Aortae were washed in PBS three times and rinsed in 100% propylene glycol, followed by staining with 0.5% Oil Red O solution in propylene glycol for 20 mins at 65°C. The samples were then put in 85% propylene glycol for 2 mins, followed by three washes in distilled water and then dried. Imaging was performed using a Nikon SMZ1500 stereomicroscope, SPOT Insight 2Mp Firewire digital camera, and SPOT Software 5.1.

### RNA Isolation and Quantitative Real-time PCR

Total RNA was extracted from primary MAECs using RNeasy kits (Qiagen)^24^. On-column DNA digestion (Qiagen) was carried out during RNA extraction. For the synthesis of the first strand of cDNA, 1 μg of total RNA after DNase treatment was reverse transcribed using cDNA Synthesis Kit (Vazyme). Quantitative real-time PCR was performed on a CFX96 detection system (StepOnePlus Real-Time PCR Systems) with SYBR Green reagents (Vazyme). All PCR primer sequences are presented in the Supplemental Tables.

### Evaluation of EndoMT Markers by Immunostaining *In Vivo* and *In Vitro*

Samples from ApoE^−/−^ and EC-iCD45KO/ ApoE^−/−^ mice were evaluated by co-staining CD31 (or VE-Cadherin) with α-SMA, KLF2, and ICAM-1. EndoMT is expressed by the overlapping percentile of CD31 (or VE-Cadherin) with other markers. For *in vitro* EndoMT marker staining, mouse aortic EC (MAEC) cultures were treated with oxLDL (100 µg/mL) or TGFβ1 (10 ng/mL) or TGFβ1 (5ng/mL) on the indicated days.

### Immunofluorescence Staining

#### Human Tissue Paraffin Sections

Human healthy control and diseased aortic arch samples from atherosclerosis patients were purchased from Maine Medical Center Biobank (Sample ID: RA06-1040A1)^24^. The paraffin sections were de-paraffinized and subjected to antigen retrieval to unmask antigenic epitopes with 10 mM Sodium Citrate, pH 6.0, and 0.5% Tween 20 at 90 °C for 30 mins^23^. Immunofluorescence staining of slides was performed using the protocol described below. Samples were blocked in PBS with donkey serum, 3% BSA, and 0.3% Triton X-100 at 4°C overnight and incubated with primary antibodies VE-Cadherin with CD45 at 4°C overnight^40^. Sections were washed three times, and respective secondary antibodies conjugated to fluorescent labels (Alexa Flour 594, 488, or 647) were added for 2 hrs at room temperature. Sections were mounted with Fluoroshield (R&D Systems) mounting medium containing DAPI (1:100).

#### Mouse aortic root and BCA cryosections

Sections were fixed with 4% paraformaldehyde for 15 min at room temperature and blocked in PBS solution containing 3% donkey and/or 10% goat serum, 3% BSA, and 0.3% Triton X-100 for 1 hr^23,39^. Samples were then incubated with primary antibody at 4°C overnight, followed by incubation with respective secondary antibodies conjugated to fluorescent labels (Alexa Flour 594, 488, or 647; 1:200) for 2 hrs at room temperature. Sections were mounted with Fluoroshield (R&D Systems) mounting medium containing DAPI (1:100).

### *En Face* Preparation and Immunofluorescent Labeling

*En face* immunofluorescence was performed as described^41^. Thoracic aortae were isolated and fixed in 2% paraformaldehyde solution at 4 °C for 1.5 hrs. Fixed tissues were permeabilized in PBS containing 0.2% Triton X-100 (permeabilization buffer) for 1 hr at room temperature and incubated with a blocking buffer (5.5% FBS in permeabilization buffer) for 1 hr at room temperature. Primary antibodies were diluted in staining buffer (2.75% FBS in permeabilization buffer) and incubated with tissue for 16 hrs at 4 °C with gentle agitation. Tissues were washed 3× in permeabilization buffer in 30-min intervals. Secondary antibodies (conjugated with Alexa Fluor 647, Alexa Fluor 594, or Alexa Fluor 488) diluted as a 1:200 working solution (10 µg/mL) in the staining buffer were added after the third wash and incubated at room temperature for 3 hrs. Tissues were washed 3× in permeabilization buffer and 1× in PBS for 30 mins at each step. After the PBS wash, vascular tissues were bisected along the direction of flow and mounted with Fluoroshield (R&D Systems). Images were obtained using a Zeiss LSM 880 microscope and Zen Black software. Zen Blue software was used for image export and analysis. Unless specified, secondary antibodies for immunohistochemistry were applied at a concentration of 1:200.

### Data Analysis

For relative gene expression data (qPCR) analysis, we use the Livak-Schmittgen method (2−ΔΔCq), which compares two values in the exponent that represent the normalized expression values for a gene of interest in samples relative to Actin or CDH5 between WT and EC-iCD45 KO cells^42^. For immunofluorescence (IF) staining, the extracted nucleus regions were split into isolated nuclei before counting using DAPI staining. Two-tailed unpaired Student’s t-tests were applied to EC-iCD45KO/ApoE^−/−^ and WT/ApoE^−/−^ atherosclerotic mice. Data are expressed as the mean ± SEM. The means of two groups were compared using Student’s t-test (unpaired, two-tailed) with P < 0.05 considered to be statistically significant. The lipid panel test was analyzed by Column Statistics and data are expressed as the mean ± SE. Unless indicated, all experiments were repeated at least three times. All data were analyzed by Prism 8 (GraphPad Software).

## RESULTS

### CD45 Expression in the Endothelium of Atherosclerotic Lesions

We first determined if CD45 was present in the endothelium of the aortic arch in pathological specimens from human patients with atherosclerosis. We detected CD45 in VE-Cadherin^+^ cells in aortic arch samples from patients with severe and mild atherosclerotic lesions (**Figure 1A**). By analyzing publicly available human scRNA-seq profiles, we observed that CD45 mRNA expression was upregulated in atherosclerosis patients compared to healthy controls **(Figure 1B**). In contrast, genes that maintain normal endothelial function such as CDH5, EMCN and FLT1 were downregulated in ECs from atherosclerotic samples (**Figure 1B**). These data suggest that upregulation of CD45 is associated with endothelial dysfunction.

**Figure 1.**
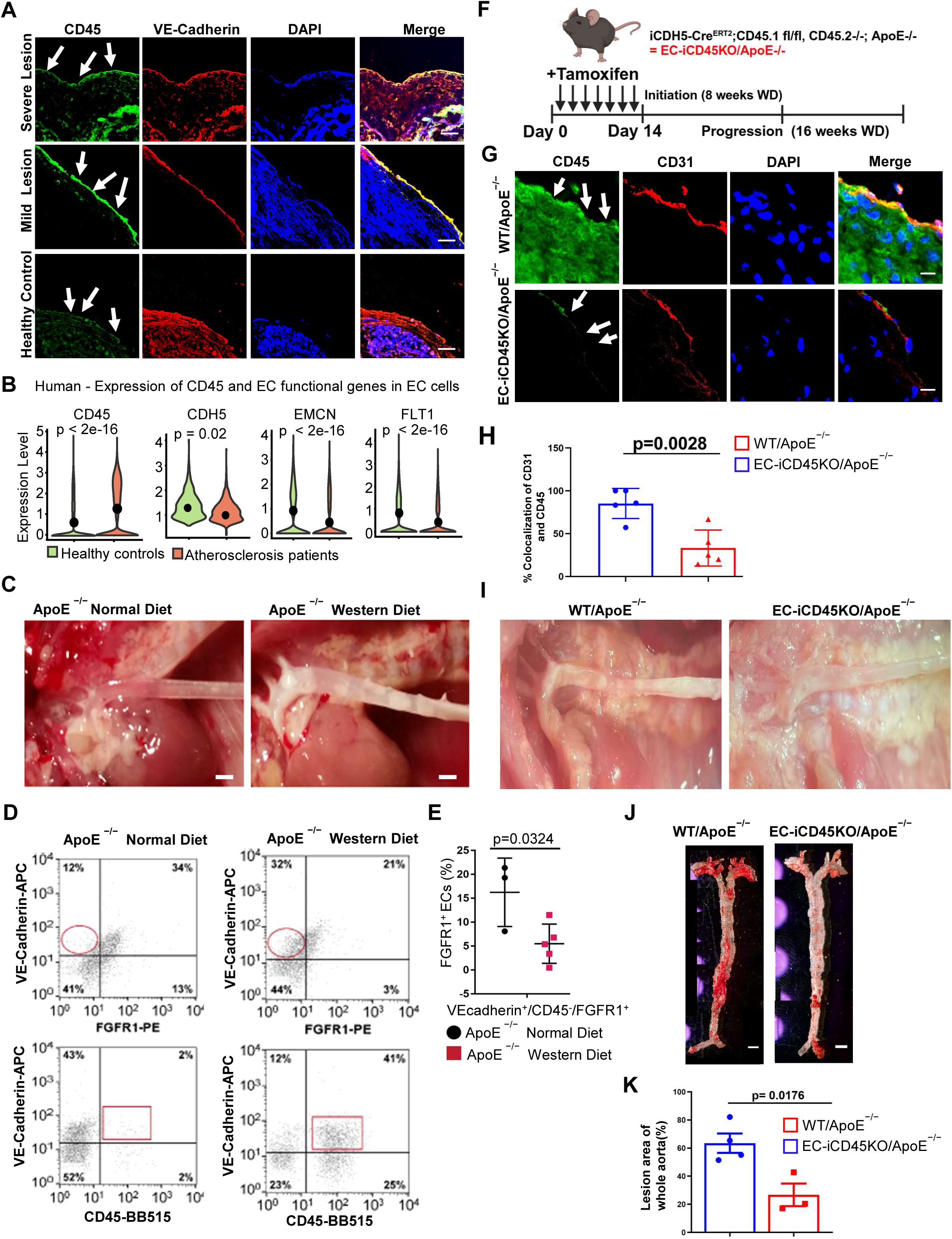
Endothelial CD45-dependent EndoMT during atherosclerosis. **A**, Immunofluorescence staining of the aortic arch section from atherosclerosis patients for CD45 (green) and VE-Cadherin (red) (Scale bar = 10 μm). **B**, Single-cell expression profiles of CD45 and endothelial functional genes in CD31 expressing cells of human healthy control and atherosclerosis patients. **C**, Aortas from ApoE^−/−^ mice fed a normal diet versus WD — lesions are apparent in ApoE^−/−^ WD-fed aortas (Scale bar = 10 μm). **D**, Representative flow cytometry analysis of ECs labeled with anti-VE-cadherin-FITC, anti-CD45-CF647, and anti-FGFR1-PE. Circles (n = 3) highlight increased FGFR1-negative ECs in ApoE^−/−^ WD aortas. Squares (n = 5) indicate increased CD45 ECs in WD-fed aortas from ApoE^−/−^ mice. **E**, Flow cytometric analysis of FGFR1+ EC from ApoE^−/−^ normal diet (black circles, n = 3) and ApoE^−/−^ WD (red squares, n = 5) mice. FGFR1+ ECs are decreased (p = 0.0324) in ApoE^−/−^ WD-fed mice aortas. **F**, Diagram showing the timeline for the deletion induced by tamoxifen injections. **G**, Immunostaining of aortic root sections from WT/ApoE^−/−^ and EC-iCD45KO/ApoE^−/−^ mice fed with western diet (WD) for 12 weeks for CD45 (green) and CD31 (red). **H**, Quantification of CD31+/CD45+ cells WT/ApoE^−/−^ and EC-iCD45KO/ApoE^−/−^ mice fed with western diet (WD) for 12 weeks. **I**, Aortas from WT/ApoE^−/−^ mice versus EC-iCD45KO/ApoE^−/−^ mice fed with western diet (WD) for 16 weeks. **J**, *En face* aortas from WT/ApoE^−/−^ and ECiCD45KO/ApoE^−/−^ female mice stained with Oil Red O (16-week WD) (Scale bar = 200 μm). **K,** Quantification of lesion area in the aortas.

As ApoE^−/−^ mice fed a normal diet also develop lesions, albeit at a slower rate compared to WD-fed ApoE^−/−^ mice, we chose to use WT mice that were fed normal chow as our true negative control for atherosclerosis occurrence to rule out the potential effect of high cholesterol content in WD-fed mice and lesion development in normal chow-fed ApoE^−/−^ mice (**Figure 1C**). When comparing aortas from ApoE^−/−^ mice fed a normal diet versus WD, lesions were apparent in ApoE^−/−^ WD-fed aortas. Flow cytometric analyses of total aortic cells for VE-Cadherin, FGFR1 and CD45 showed a significant reduction in VE-Cadherin^+^/FGFR1^+^ ECs in WD-fed ApoE^−/−^mice, compared to normal diet-fed ApoE^−/−^ mice (**Figure 1D** and **1E**). In contrast, VE-Cadherin^+^/CD45^+^ ECs were increased in WD-fed ApoE^−/−^ mice compared to normal diet-fed ApoE^−/−^ mice, suggesting loss of FGFR1 may contribute to EndoMT in lesions of WD-fed ApoE^−/−^ mice (5.5 vs 16.3%, p = 0.0324).

We generated mice with tamoxifen-inducible deletion of endothelial CD45 on an ApoE^−/−^background (EC-iCD45KO/ApoE^−/−^) (**Figure 1F** and **Figure S1A**,**B**). Mice from either genotype fed a WD diet for 16 weeks (WT/ApoE^−/−^) showed no difference in body weight for males or females (data not shown). Serum total cholesterol and triglycerides measurements (124.2.8±28.46 vs 103.6 ± 19.02; p value = 0.6700); HDL (73.27 ± 13.99 vs 79.44 ± 18.03; p value = 0.9270, LDL (728.6 ± 111.9 vs 762.4 ± 117.4; p = 0.5516), showed no significant difference between WT/ApoE^−/−^ and EC-iCD45KO/ApoE^−/−^ mice (**Figure S2**). We then analyzed expression and localization of CD45 in mouse aortic ECs by immunofluorescence staining of sections from the aortic root. CD45 was detected and co-expressed with CD31 in 12-week-old ApoE-null mice (**Figure 1G** and **1H**). Intriguingly, there was a significant decrease in CD45 in ECs from EC-iCD45KO/ApoE^−/−^ mice (∼52.05%; p = 0.0028). Aortas from WT/ApoE^−/−^ mice fed a WD showed more lesions when compared to EC-iCD45KO/ApoE^−/−^ WD-fed aortas (male, 16-week WD) (**Figure 1I**). Consistent with this observation, whole-mount *en face* aortas from female WT/ApoE^−/−^ and EC-iCD45KO/ApoE^−/−^mice stained with Oil Red O after a 16-week WD had a reduction in lesion size (∼36.78%; p = 0.0176) (**Figure 1J** and **1K**). Together, these results show that CD45 depletion significantly reduces atherosclerotic burden in ApoE-null mice.

### Single-Cell Profiling Shows CD45 Deletion Suppresses EndoMT and Reshapes Vascular Cell Identity in Atherosclerosis

To establish that endothelial CD45 plays a decisive role in atherosclerosis progression, we profiled single-cell transcriptomes of atherosclerotic ApoE^−/−^ and EC-iCD45KO/ApoE^−/−^ mice using scRNA-seq and bioinformatic analyses. WT/ApoE^−/−^ and EC-iCD45KO/ApoE^−/−^ male mice were fed a WD for 16 weeks, followed by isolation of aortas, enzymatic digestion, scRNA-seq and data analysis (**Figure S1C**). Through stringent data preprocessing, we retained high-quality cells by filtering based on the number of genes expressed, doublets as well as mitochondrial and ribosomal gene expression signatures (**Figure S3A**). We also integrated single-cell datasets from WT/ApoE^−/−^ and EC-iCD45KO/ApoE^−/−^ mice, corrected for batch effects and subsequently performed single-cell clustering and downstream analyses. We confirmed the efficacy of CD45 knockout in aortic ECs (**Figure S3B**) using single-cell transcriptomic profiles.

Through our single-cell data analysis, we identified 25 distinct clusters, which were grouped into major cell types (**Figure S3C**, left), including vascular cells (*e.g.*, ECs and VSMCs) and immune cells (*e.g.*, macrophages, B cells, T cells and others). We then focused on vascular cells to investigate the association of CD45 with the EndoMT process (**Figure S3C**, right). Within this subset, we identified five vascular cell types based on their marker gene expression signatures (**Figure 2A**): ECs (Pecam1, Cdh5), EndoMT1 cells (Fbln5, Fbln2), EndoMT2 cells (Tgfb1, Tgfbr2), VSMCs (Acta2, Tagln), and proliferating cells (Top2a, Mki67) (**Figure S3D**).

**Figure 2.**
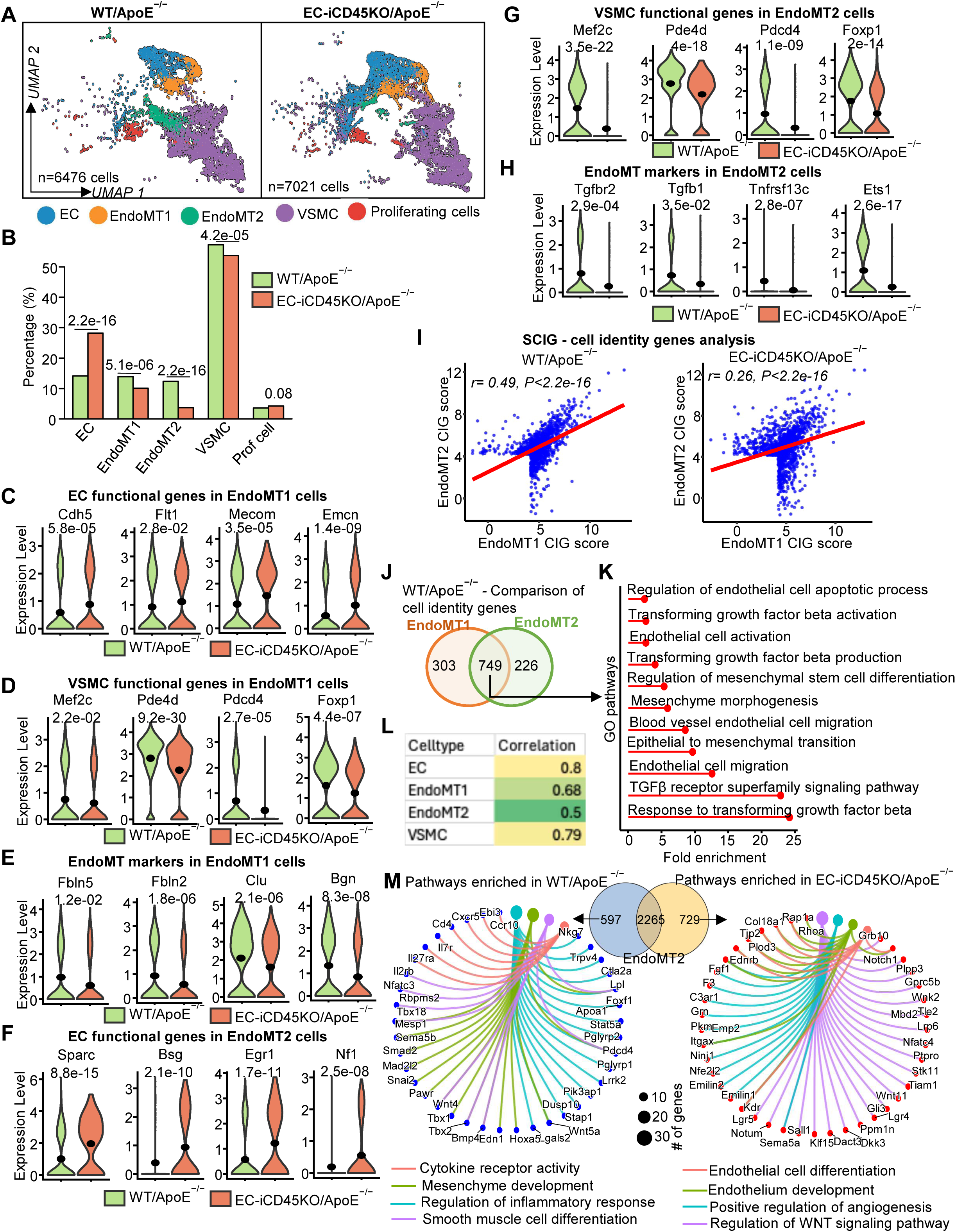
Endothelial cell-specific deletion of CD45 rewires vascular cell identity in WT/ApoE^−/−^ and EC-iCD45KO/ApoE^−/−^ mice. **A**, Five vascular cell types were identified using scRNA-seq data from WT/ApoE^−/−^ and EC-iCD45KO/ApoE^−/−^ mice. **B**, Bar graph depicting the percentage of each cell population in aortas from WT/ApoE^−/−^ and EC-iCD45KO/ApoE^−/−^ mice. **C-H**, Expression levels of endothelial (C, F), VSMC (D, G) functional genes and EndoMT markers (E, H), shown by violin plot, in EndoMT1 (C-E) and EndoMT2 (F-H) cells between WT/ApoE^−/−^ and EC-iCD45KO/ApoE^−/−^ mice. **I**, Scatter plot showing the Pearson correlation between cell identity gene (CIG) scores of EndoMT1 and EndoMT2 cells in WT/ApoE^−/−^ (left) and EC-iCD45KO/ApoE^−/−^ (right) mice. **J**, Venn diagram depicting shared and unique cell identity genes identified by the SCIG tool in EndoMT1 and EndoMT2 cells of WT/ApoE^−/−^ mice. **K**, Pathway analysis of shared CIGs EndoMT genes between EndoMT1 and 2 cells strongly implicated in EndoMT processes. **L**, Pearson correlation of cell identity gene scores for each vascular cell type between WT/ApoE^−/−^ and EC-iCD45KO/ApoE^−/−^ mice. **M**, Gene concept diagrams illustrate Gene Ontology (GO) pathway enrichment of unique cell-identity genes identified in EndoMT2 cells. In WT/ApoE^−/−^ mice (left), enriched pathways are primarily associated with inflammatory, cytokine-mediated, and muscle-related functional processes. In EC-iCD45KO/ApoE^−/−^ mice (right), enriched pathways are predominantly related to endothelial cell function and angiogenesis.

Comparative analysis of cell populations in WT/ApoE^−/−^ and EC-iCD45KO/ApoE^−/−^ aortas revealed a significant increase in the EC population (from 14% to 28%) and a reduction in vascular SMCs (from 58% to 53%) in EC-iCD45KO/ApoE^−/−^ mice compared to ApoE^−/−^ mice (**Figure 2B**). We characterized EndoMT cells into two subclusters—EndoMT1 and EndoMT2 (**Figure 2A**). In the EndoMT cell clusters, the population of EndoMT1 (∼3%) and EndoMT2 (∼9%) significantly decreased in the EC-iCD45KO group (**Figure 2B**). These findings suggest that the loss of CD45 in ECs contributes to rewiring of vascular cell identity and function in atherosclerotic ApoE^−/−^ mice.

Next, we compared the gene expression signatures in EndoMT cell types between two groups (*i.e.*, EC-iCD45KO/ApoE^−/−^ and WT/ApoE^−/−^ mice). Violin plots showed that EC function genes (Cdh5, Flt1, Mecom) were upregulated in EC-iCD45KO/ApoE^−/−^ aortas compared with WT/ApoE^−/−^ aortas (**Figure 2C**). At the same time, SMC function genes (Mef2c, pde4d, pdcd4) and EndoMT markers (Fbln2, Fbln5, Clu) were downregulated in EC-iCD45KO/ApoE^−/−^ mice in EndoMT1 cells (**Figure 2D** and **2E**). Similarly, in EndoMT2 cells, EC function genes (Sparc, Bsg, Egr1) increased (**Figure 2F**), but SMC function genes (Mef2c, pde4d, pdcd4) and EndoMT markers (TGFβ1, TGFβR2, Ebf1) decreased in EC-iCD45KO/ApoE^−/−^ mice when compared to WT/ApoE^−/−^ mice (**Figure 2G** and **2H**).

### SCIG Analysis Illustrates CD45-Deficiency Reprograms EndoMT Cells Toward an Endothelial Identity while Suppressing Mesenchymal and Inflammatory Signatures

Recently, we showed that SCIG^32^ effectively profiles the full spectrum of cell identity genes (CIGs) that enables detailed characterization of individual cell types and their functional states in atherosclerotic and control samples^43^. To characterize transcriptomic and functional differences between EndoMT1 and EndoMT2 cells, we identified CIGs using SCIG and performed GO pathway analysis. Unlike differential expression analysis, which compares gene expression between conditions, the SCIG algorithm is designed to uncover CIGs within a given cell type by integrating gene expression values with gene sequence signatures. Genes with an FDR < 0.05 in SCIG predictions were considered potential CIGs. Pearson correlation analysis of CIGs and their scores revealed that EndoMT1 and EndoMT2 cells share substantial cell identity, with a correlation of 0.49 in WT/ApoE^−/−^ mice compared to a reduced correlation of 0.26 in EC-iCD45KO/ApoE^−/−^ mice (**Figure 2I**). These findings suggest that EndoMT2 cells underwent the most pronounced transcriptional and cellular changes between the two groups.

We further observed that 749 CIGs were shared between EndoMT1 and EndoMT2 cell types (**Figure 2J)**, many of which were implicated in EndoMT-related processes such as EC migration, activation of TGF-β signaling and mesenchymal cell development (**Figure 2K**). Pathway analysis of uniquely enriched CIGs (226 genes) revealed that EndoMT2 cells were functionally associated with immune processes and inflammatory signaling, including B-, T-, and mast cell activation, mononuclear cell proliferation, interleukin production, muscle cell development and inflammatory responses (**Figure S4A**, right). In contrast, unique CIGs in EndoMT1 cells (303 genes) were enriched in pathways primarily associated with inflammatory responses, mesenchymal cell development and EC function (**Figure S4A**, left). Moreover, we observed that CD45 is highly expressed in both EndoMT1 and EndoMT2 cells of WT/ApoE^−/−^mice, suggesting a role for CD45 in regulating EndoMT during atherosclerosis (**Figure S3B**).

To investigate the role of CD45 in determining cell identity, we applied the SCIG algorithm to each vascular cell type by identifying CIGs in WT/ApoE^−/−^ and EC-iCD45KO mice. Pearson correlation analysis of CIG scores between the two groups revealed the lowest correlation for EndoMT2 cells (r = 0.50), followed by EndoMT1 (r = 0.68), VSMCs (r = 0.79) and ECs (r = 0.80) (**Figures 2L** and **S4B**). These results indicate the cell identity of EndoMT1 and EndoMT2 populations is more divergent and modulated by CD45 status compared to VSMCs and ECs.

Based on these findings, we performed pathway analysis on the uniquely identified CIGs in WT/ApoE^−/−^ and EC-iCD45KO mice for both EndoMT1 and EndoMT2 cells (**Figures 2M**, and **S5A** to **S5G**). Pathways related to mesenchymal development and differentiation, epithelial cell fate, SMC contraction and proliferation as well as cytokine and inflammatory responses were significantly enriched among uniquely identified CIGs in EndoMT1 and EndoMT2 cells from WT/ApoE^−/−^ mice (**Figures S5B** and **S5F**), but not in EC-iCD45KO mice. The endothelial function pathways, including EC proliferation, vascular development, VEGF and WNT signaling and angiogenesis were enriched among uniquely identified CIGs in both EndoMT1 and EndoMT2 cells from EC-iCD45KO mice (**Figures S5D** and **S5G**).

Additionally, we visualized these pathways through gene concept network diagrams that depict pathway nodes and their connecting gene edges to highlight functional relationships (**Figures 2M**, **S5C** and **S5E**). For instance, in WT/ApoE^−/−^ mice, EndoMT2 cells showed enrichment of mesenchymal development pathways driven by genes such as Snai2, Smad2 and Pdcd4, along with cytokine receptor activity genes including Il7r, Il2rb, Cxcr5 and Ccr10 (**Figure 2M**, left). In contrast, EC-iCD45KO mice exhibited enrichment of endothelium development and differentiation pathways in EndoMT2 cells with genes such as Kdr, Notch1, Tie2, Tjp2 and Ednrb supporting endothelial function programs (**Figure 2M**, right). Together, these identity gene signatures, and pathway analyses reveal CD45 loss attenuates mesenchymal-like transition programs and promotes endothelial identity and function.

### Single-Cell Dynamics Analyses Shows Deletion of Endothelial CD45 Inhibits EndoMT in Atherosclerosis

To further study the role of CD45 in EndoMT, we performed single-cell dynamics analysis, which enables the study of how individual cells differentiate into other cells. First, we applied RNA velocity analysis to the cells from each group and found that ECs in WT/ApoE^−/−^ mice exhibited strong transitions toward EndoMT1 and EndoMT2 populations, eventually acquiring a VSMC phenotype (**Figure 3A**, left). In contrast, this transition was markedly reduced in EC-iCD45KO/ApoE^−/−^ mice (**Figure 3A**, right). Trajectory analysis using Monocle3 revealed that in WT/ApoE^−/−^ mice, the inferred trajectory progressed from ECs along two distinct paths, EndoMT1 and EndoMT2, ultimately leading toward VSMCs (**Figure 3B**) when ECs were selected as the root cells. Along this trajectory in WT/ApoE^−/−^ mice, endothelial function genes (Pecam1, Vwf, Kdr) were downregulated, indicating a loss of EC identity, while upregulation of VSMC function genes (Myl9, Acta2, Tagln) suggested a shift toward a VSMC-like phenotype (**Figure 3C** and **3D**).

**Figure 3.**
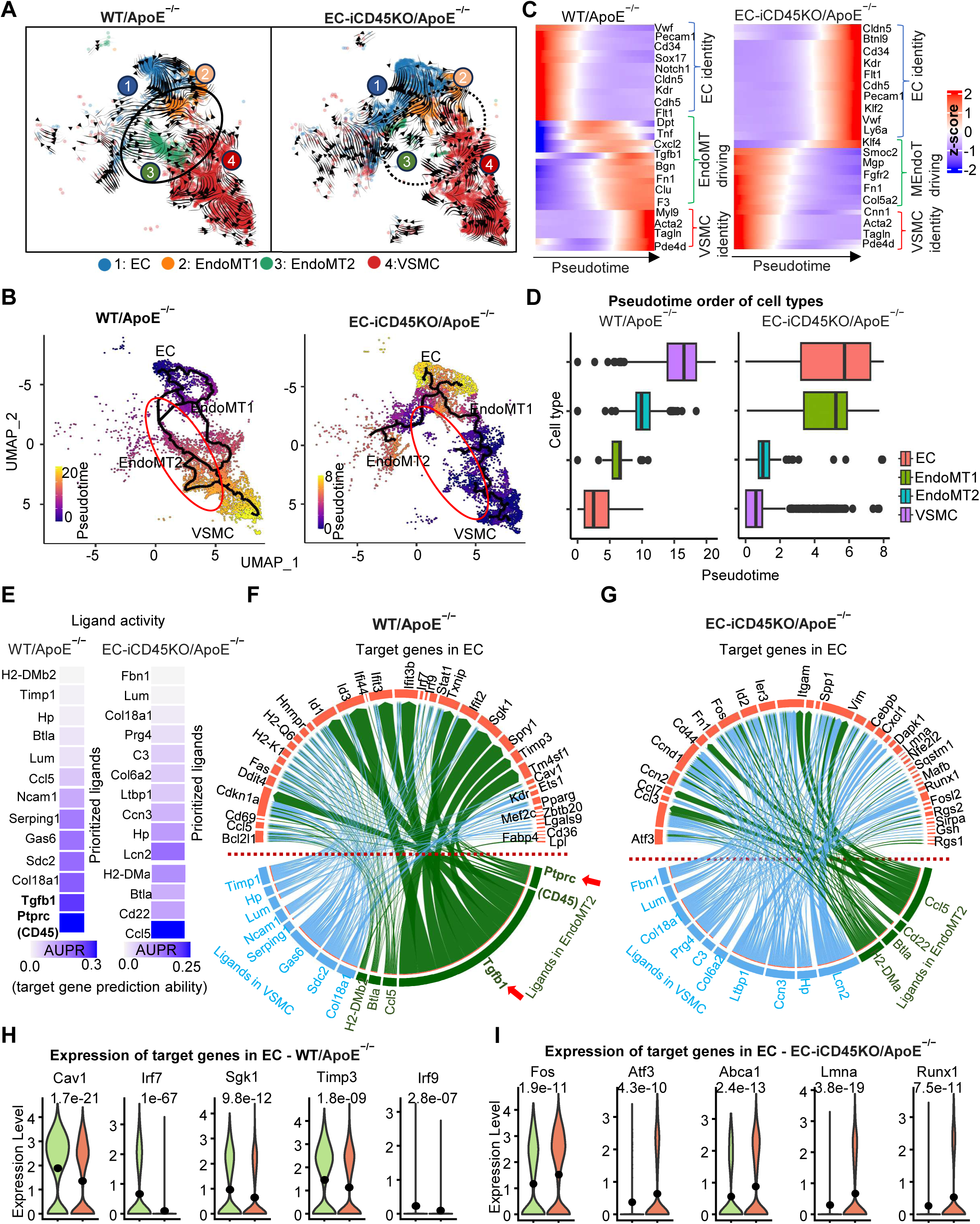
Role of CD45 in EndoMT using single-cell dynamics and cell-cell communication analyses. **A**, RNA velocity analysis shows the transition of endothelial cells (ECs) to vascular smooth muscle cells (VSMCs) through EndoMT cells in WT/ApoE^−/−^ mice (left), a process that is absent in EC-iCD45KO/ApoE^−/−^ mice. Instead, a mesenchymal-to-endothelial transition (MEndoT) process is observed (right). Solid black outlines indicate the EndoMT trajectory, while dotted outlines indicate the MEndoT trajectory. **B**, Monocle3 trajectory inference analysis comparing WT/ApoE^−/−^ and EC-iCD45KO/ApoE^−/−^ mice. **C**, Gene expression changes associated with the EndoMT and MendoT trajectories in WT/ApoE^−/−^ and EC-iCD45KO/ApoE^−/−^ mice, respectively. **D**, Boxplots showing the ranking of vascular cell types based on their pseudotime in WT/ApoE^−/−^ (left) and EC-iCD45KO/ApoE^−/−^ (right) mice. **E**, Potential ligands identified in sender cells of WT/ApoE^−/−^ and EC-iCD45KO/ApoE^−/−^ mice. **F-G**, Chord diagrams illustrating potential ligands and their regulation of target genes in receiver ECs of WT/ApoE^−/−^ mice (F) and CD45-deletion mice (G). **H-I**, Expression profiles of identified target genes in receiver ECs of WT/ApoE^−/−^ mice (H) and CD45-deletion mice (I)

EndoMT1 and EndoMT2 cells represent intermediate states, characterized by upregulation of EndoMT markers (Tgfb1, Bgn, Clu) and inflammatory markers (Tnf, Cxcl2), bridging ECs and VSMCs (**Figure 3C, left)**. In addition, when EndoMT1 or EndoMT2 cells were used as the root, the trajectories also terminated in VSMCs (**Figure S6A** and **S6B**). VSMC function gene expression patterns aligned with the EndoMT process trajectory (**Figure S6C** and **S6D**) in WT/ApoE^−/−^ mice. However, through unbiased RNA velocity analysis, the EndoMT transition was markedly inhibited in EC-iCD45KO mice where VSMCs tended to revert toward an EC-like state (**Figure 3A, right**). Consistent with the RNA velocity results, the inferred trajectory (**Figures 3B, right** and **3D right**) and associated gene expression changes (**Figure 3C, right**) support a VSMC-to-EC transition (MEndoT, Mesenchymal-to-Endothelial Transition). Results from both RNA velocity and trajectory analyses were consistent, demonstrating that EndoMT was prevalent in WT/ApoE^−/−^ mice but significantly suppressed and reversed in EC-iCD45KO mice.

### CD45 Drives EndoMT Through Cell-Cell Communication Signals

To identify potential cell-cell communication signals in the EndoMT process, we used the NicheNet tool to prioritize ligands and their target genes based on single-cell expression profiles. In this analysis, EndoMT2 and VSMC cells were designated as sender cells, while ECs served as receiver cells (**Figures 3E** to **3I**). Using the NicheNet pipeline, we identified potential ligands in EndoMT1 cells of WT/ApoE^−/−^ mice, including Ptprc/CD45, Tgfβ1, and in VSMCs (**Figure 3E**, left), including Col18a1, Lum, and Timp1 (**Figure 3E**, left). These ligands were found to regulate target genes such as Irf7, Cav1, Sgk1, and Timp3, which are involved in accelerating EndoMT (**Figure 3F**). Expression levels of the identified target genes were decreased in CD45-deletion mice (**Figure 3H)**. Notably, in CD45-deletion mice, these ligand-target gene signaling interactions were diminished (**Figure 3G)**. We observed the target genes Fos, Atf3, Abca1, Lmna and Runx1 that are involved in endothelial identity, proliferation and cholesterol efflux, were upregulated in CD45-deletion mice (**Figure 3I**).

### Endothelial CD45-Deficiency Inhibits EndoMT by Preventing TGFβR Signaling and Increasing FGFR2 Expression during Atherosclerosis

We performed *en face* immunofluorescence staining of the thoracic aorta with antibodies to VE-Cadherin, CD31, DAPI and CD45. This showed CD45 co-expression with VE-Cadherin and CD31 in WT ApoE-null mice but a significant reduction of CD45 expression (22.95%; p = 0.0308) in EC-iCD45KO ApoE-null mice (**Figure 4A** to **4C**). Endothelial dysfunction is a critical event in vascular inflammation that is characterized, in part, by elevated cell surface expression of adhesion molecules such as intercellular adhesion molecule-1 (ICAM-1) and vascular adhesion molecule-1 (VCAM-1). Accordingly, we observed a reduction of ICAM-1 expression (25.22%;p = 0.0087) in ECs from the thoracic aortas of EC-iCD45KO/ApoE^−/−^ mice and a similar reduction in aortic roots from these mice (**Figure 4D** to **4F** and **S7A**) compared to WT/ApoE^−/−^ mice. In contrast, VCAM1 expression in the aortic root did not show a significant difference between WT/ApoE^−/−^ and EC-iCD45KO/ApoE^−/−^ mice (**Figure S7B**).

**Figure 4.**
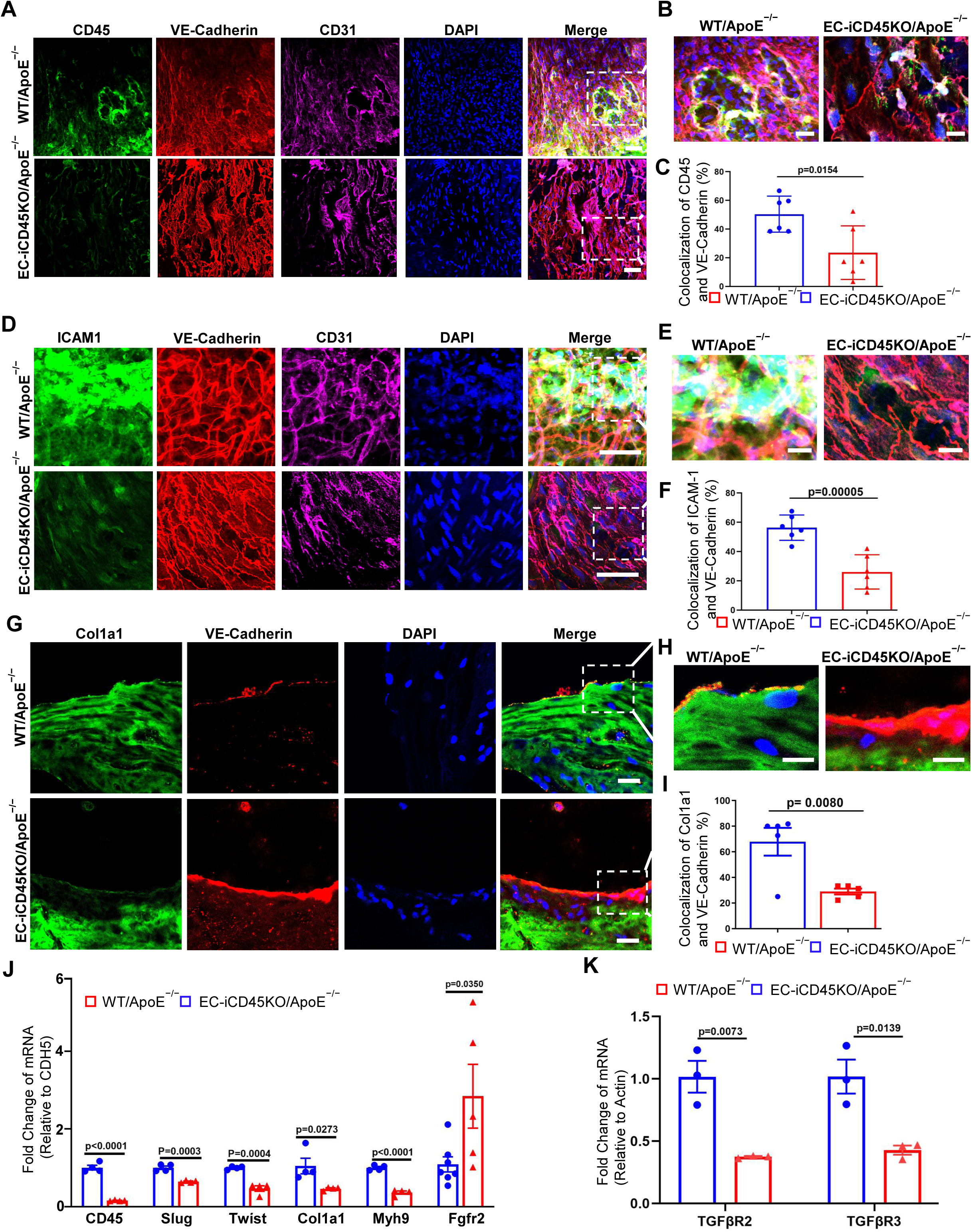
Endothelial CD45-deficiency inhibits EndoMT by reducing TGFβ signaling while upregulating FGFR2 expression during atherosclerosis. **A**, *En face* immunostaining of the thoracic aorta was performed with antibodies to CD45, VE-Cadherin, and CD31; DAPI staining shown to visualize nuclei (Scale bars = 50 μm). **B**, Enlarged images of immunostaining for CD45 and CD31 and from Figure 4A (Scale bars = 10 μm). **C**, Quantification for colocalization of VE-Cadherin and CD45. **D**, *En face* immunostaining of the thoracic aorta was performed with antibodies to ICAM-1, VE-Cadherin, and CD31; DAPI staining shown to visualize nuclei (Scale bars = 50 μm). **E**, Enlarged images of immunostaining for ICAM-1 and CD31 from panel D (Scale bars = 20 μm). **F**, Quantification of colocalized VE-Cadherin and ICAM-1 in the thoracic aorta. **G**, Aortic root sections from WT/ApoE^−/−^ and EC-iCD45KO ApoE^−/−^ mice were co-stained with the Col1a1 and VE-Cadherin. **H**, Enlarged images of immunostaining for Col1a1and VE-Cadherin from Figure 4 **G** (Scale bars = 20 μm). **I**, Quantification of colocalized Col1a1 and VE-Cadherin in the aortic root. **J**, MAECs isolated from of WT/ApoE^−/−^ and EC-iCD45KO/ApoE^−/−^ mice were treated with 5 µmol/L tamoxifen for 3 days, followed by treatment with 100 µg/mL oxidized low-density lipoprotein, and then lysed for quantitative polymerase chain reaction to detect endothelial-to-mesenchymal transition (EndoMT) markers (n = 3). Reductions in expression of Twist and Slug, genes associated with the EndoMT process, were seen in CD45 KO MAECs. MYH9 and Col1a1 were also significantly reduced, while FGFR2 was significantly upregulated. **K**, TGFβR2 and TGFβR3 were significantly reduced in endothelial CD45 KO MAEC.

Because collagen type I alpha 1 (Col1a1) is a crucial structural component of the fibrous cap in atherosclerotic plaques, we assessed the expression of this protein in aortic roots. We found a reduction of Col1a1 immunofluorescence staining in EC-iCD45KO/ApoE^−/−^ mice (**Figure 4 G**). We also determined that Col1a1 colocalized less with VE-cadherin in EC-iCD45KO/ApoE^−/−^ mice when compared to WT/ApoE^−/−^ mice (38.88%; p = 0.0080) (**Figure 4H** and **4I**). This indicated a reduction of collagen content in ECs in EC-iCD45KO/ApoE^−/−^ mice. In addition, we detected upregulation of KLF2 (1.717-fold; p = 0.0175) in the aortic roots from EC-iCD45KO/ApoE^−/−^ mice fed a WD for 12 weeks when compared to WT/ApoE^−/−^ mice (**Figure S8A** and **S8B**). These findings are consistent with our observation that EndoMT cell clusters were functionally associated with immune processes and inflammatory signaling.

Our results indicate that loss of endothelial CD45 results in diminished atherosclerosis while TGFβ signaling and EndoMT contribute to atherosclerosis. To model this *in vitro*, mouse aortic ECs (MAECs) from WT/ApoE^−/−^ and EC-iCD45KO/ApoE^−/−^ mice were isolated and treated with tamoxifen, followed by stimulation with 100 µg/mL oxLDL to initiate EndoMT. Cell mRNA was collected for quantitative RT-PCR assays to detect expression of the EndoMT markers TGFβR2 and TGFβR3. MAECs with tamoxifen-induced CD45 deletions had reduced TGFβR_­_(∼64.26%; p = 0.0073) and TGFβR_­_(∼59.04%; p = 0.0139) mRNA levels (**Figure 4J** and **4K**). We confirmed an 84.52% (p<0.0001) reduction of CD45 mRNA expression in EC-iCD45KO MAECs compared with the WT/ApoE^−/−^ control group when treated with oxLDL (100 µg/mL). Subsequent analyses showed reductions of MYH9 (∼63.44%; p<0.0001), Col1A1 (∼58.26%; p = 0.0273), Twist (∼53.15%; p = 0.0004) and Slug (∼35.58%; p = 0.0003) mRNA expression in EC-iCD45DKO MAECs compared to the control group. We also observed an upregulation of approximately 1.75-fold (p = 0.0350) for FGFR2 mRNA levels in MAECs from EC-iCD45DKO mice when compared to the ECs treated only with oxLDL. These *in vitro* results confirm that loss of endothelial CD45 decreases EndoMT marker expression and TGFβ signaling. CD45 deletion from MAECs also increases FGFR2 mRNA expression. As discussed previously, increases in FGF receptor expression and KLF2 are known to inhibit TGFβ signaling in ECs to attenuate EndoMT.

### Endothelial Loss of CD45 Prevents Inflammation and Atherosclerosis

To substantiate our findings that the loss of CD45 expression can constrain the progression of atherosclerosis, we performed histological staining on male and female ApoE^−/−^ and EC-iCD45KO/ApoE^−/−^ mice fed a WD. We evaluated the aortic sinus and abdominal aorta as well as performing *en face* preparations of the whole aorta. Whole mount *en face* aortas from ApoE^−/−^ and EC-iCD45KO/ApoE^−/−^ mice were stained with Oil Red O (**Figure 5A, B**: 8–13-week WD; **Figure 5C, 5D**: 10–13-week WD). We found that EC-iCD45KO/ApoE^−/−^ mice fed an atherosclerotic WD exhibited significant reductions in lesional area throughout the aorta (∼23.88%; ∼8-13 week WD;∼36.79%, 10-13 week WD) when compared with ApoE-null mice that expressed CD45. Overall, these results indicate that loss of endothelial CD45 prevents atherosclerosis progression.

**Figure 5.**
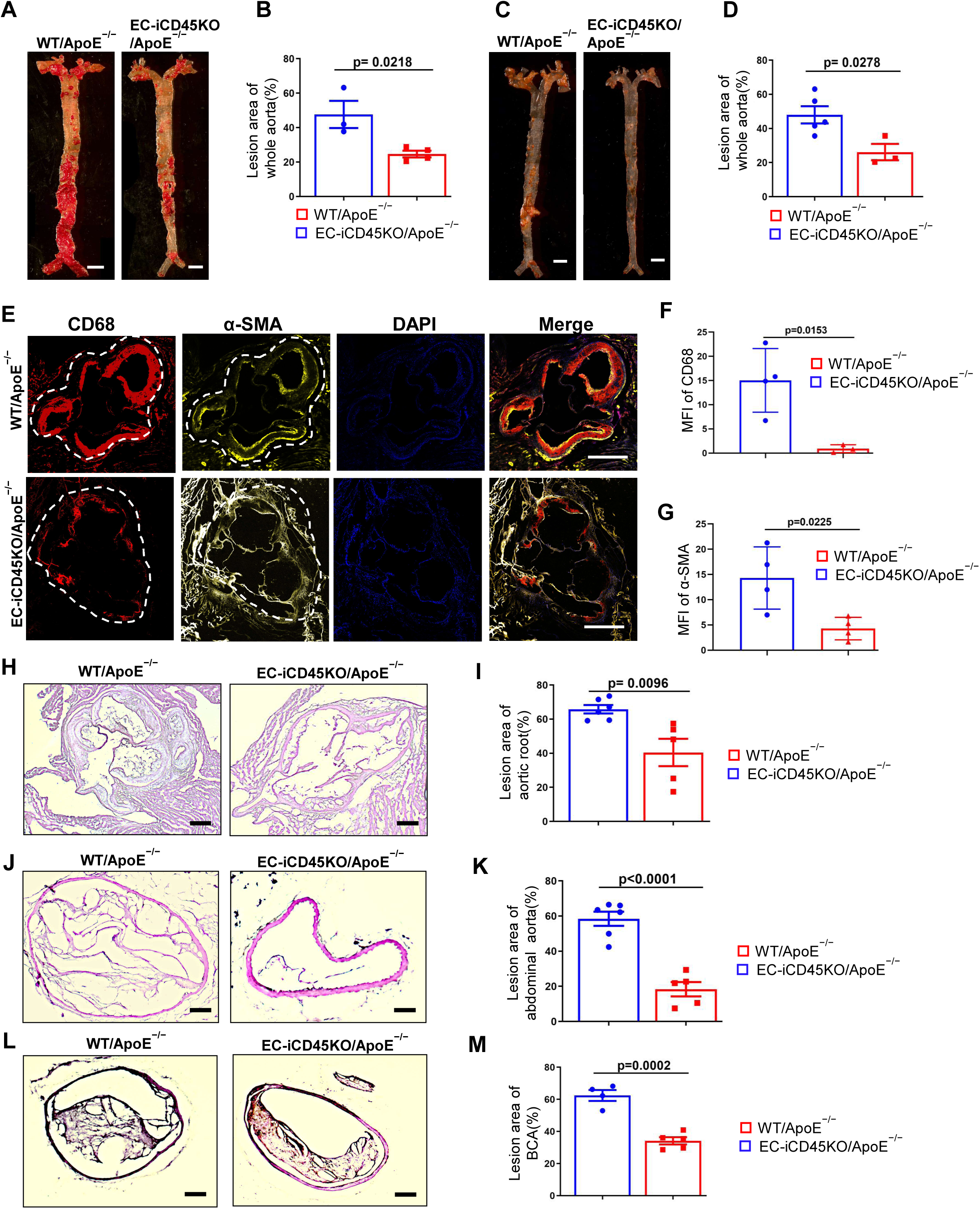
Endothelial CD45 deletion reduces atherosclerosis in ApoE^−/−^ mice. **A-D**, *En face* aorta preparations from WT/ApoE^−/−^ and EC-iCD45KO /ApoE^−/−^ male (A, 8–13-week WD) and female (C, 10-13 week WD) mice stained with Oil Red O. Quantifications for *en face* Oil Red O staining in male (B) and female mice (D) (Scale bar = 200 μm). **E-G**, Aortic root sections from WT/ApoE^−/−^ and EC-iCD45KO ApoE^−/−^ mice were stained with antibodies against the macrophage marker CD68 and α-SMA (E, Scale bar = 200 μm). Quantifications for CD68 (F) and α-SMA (G) staining. **H-M**, Hematoxylin and eosin (H&E) staining of the aortic root (H); aortic aneurysm sections (J); and brachiocephalic artery (L). Quantifications for H&E staining in the aortic root (I), aortic aneurysm sections (K), and brachiocephalic artery (M) (Scale bars = 10 μm).

To determine whether this phenomenon extends to regions of the vasculature beyond the aorta, we assessed the aortic root and BCA from ApoE^−/−^ and EC-iCD45KO/ApoE^−/−^ mice fed a WD for 16 weeks. Sections from these tissues were immunofluorescently-stained for both the macrophage marker CD68 and α-SMA. We also created a tamoxifen-inducible endothelial-specific lineage tracing system in the presence or absence of endothelial CD45 (**Figure S9A**). Aortic root sections from ApoE^−/−^/TdTomato and EC-iCD45KO/ApoE^−/−^/TdTomato mice fed a WD for 16 weeks were also used for immunofluorescence staining. We showed that α-SMA expression was reduced in the aortic root endothelium in EC-iCD45KO/ApoE^−/−^/TdTomato mice compared to ApoE^−/−^ mice (**Figure S9B**), which supports the contention that loss of endothelial CD45 inhibits EndoMT.

Additional substantiation of this assertion comes from experiments demonstrating diminished foam cell formation and α-SMA expression in arteries from CD45-deficient mice (**Figure 5E** to **5G**). These studies revealed reductions in atherosclerotic lesion prevalence in aortic roots (∼25.39%; 12-week WD), BCAs (∼28.28%; 16-week WD) and abdominal aortas (∼40.05%; 16-week WD) in endothelial CD45-deficient mice (**Figure 5H** to **5M**) using H&E staining. We also performed Oil Red O staining on aortic roots and BCAs to confirm the observed reductions in atherosclerotic lesion burden as a result of CD45 loss (**Figure S10A** to **S10D**). CD45^+^/VE-Cadherin^+^/α-SMA^+^ cells were significantly increased from 0.8 ± 1.0% in WT mice (n = 3) to 9.2 ± 0.7% in ApoE^−/−^ WD-fed mice (n = 3) (p = 0.0003) compared to normal diet-fed ApoE^−/−^mice. The presence of α-SMA^+^ in ApoE^−/−^ WD-fed mice indicates CD45^+^ ECs underwent EndoMT within an aortic atherosclerotic lesion (**Figure S10E**). This finding is consistent with the above cytometric analyses of total aortic cells for VE-Cadherin, FGFR1 and CD45 expression that showed a significant reduction in VE-Cadherin^+^/FGFR1^+^ ECs in WD-fed ApoE^−/−^ mice compared to normal diet-fed ApoE^−/−^ mice. It is also worth noting that all immunofluorescence staining described above was performed alongside experimental controls using serial sections stained with isotype-matched control IgGs (**Figure S11A to S11C**) to ensure the validity of antibody staining. A graphical depiction of our findings (**Figure 6**) highlights the novel role of CD45 in atherosclerosis progression through EndoMT regulation.

**Figure 6.**
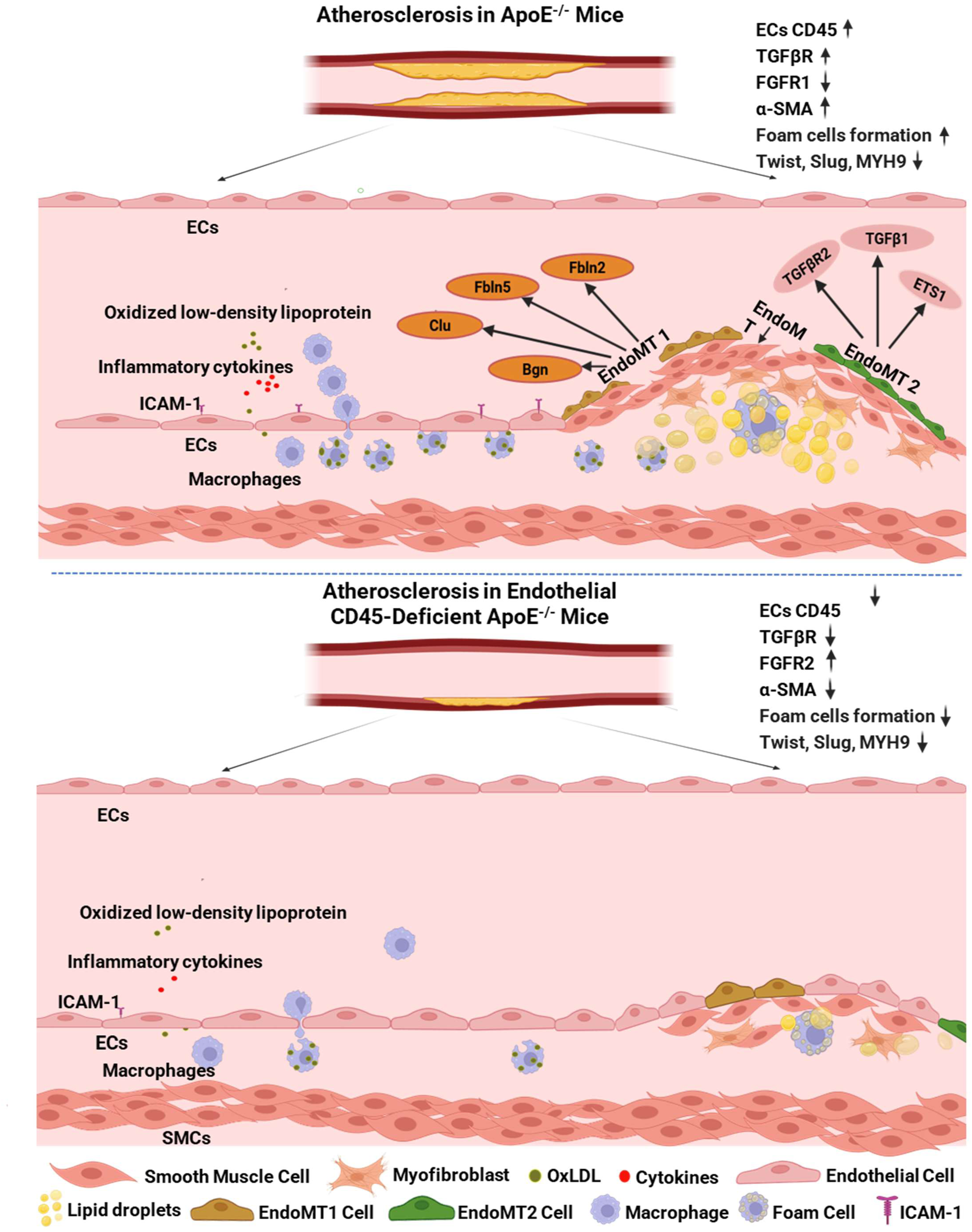
Loss of endothelial CD45 reduces EndoMT and atherosclerosis in WT/ApoE^−/−^ mice. EndoMT can be induced by several factors including TGFβ signaling, inflammatory cytokines, and expression of the protein tyrosine phosphatase CD45. During this process, CD45-positive (termed as ‘partial EndoMT’) cells contribute to neointima formation and vascular wall remodeling. Twist, MYH9, Col1a1 expression has been indicated to be reduced in EC-iCD45KO cells compared to that in the control MAECs. Expression of FGFR2 has been raised in EC-iCD45KO cells compared to that in the control mice. Additionally, it has been reported that ERK pathway levels and p-Smad2 expression were higher in the CD45-positive cells than in the cells from control mice.

## DISCUSSION

Our data show that deletion of CD45 from endothelial cells reduces atheroma burden in ApoE-null mice fed a Western diet (WD). Our single-cell experiments further indicate that EC-specific knock-out of CD45 moderates a novel sub-population of EndoMT cells, designated EndoMT2. We present evidence that CD45 deficiency leads to decreases in TGFβR_­_, TGFβR_­_, Twist, Slug, MYH9 and Col1a1 expression in MAECs pre-treated with oxLDL. In addition, we unexpectedly discovered that the absence of endothelial CD45 led to the reduction of α-smooth muscle actin and Col1a1, which are typical of EndoMT, in the aortic plaque. Finally, we found endothelial CD45 deficiency increased KLF2 expression in the aortic plaque, a transcription factor known to counter EndoMT, as well as augmenting FGFR2 expression.

Because the canonical hematopoietic marker CD45 is commonly used as a negative marker for ECs, the existence of CD45-positive ECs may have been overlooked in previous studies despite increasing evidence that indicates CD45-positive ECs are associated with certain pathological conditions^44^. In a rat pulmonary arterial hypertension model, lung ECs were found to express CD45 and endothelial function was verified by tubule formation in cocultures with lung fibroblasts^2^. Protein tyrosine phosphatase receptor type C (CD45)-positive endothelium has also been observed in the mitral valves of adult sheep 6 months after myocardial infarction resulting in an EndoMT phenotype^4^. Moreover, the CD45 phosphatase inhibitor is reported to block EndoMT activity in TGFβ1-treated ovine mitral valve ECs^4^. Taken together these studies implicate endothelial CD45 in heart valve development and cardiovascular disease.

In this study, we discovered CD45-positive ECs in the aortic arches of patients with atherosclerosis, which reinforces the notion that endothelial CD45 plays a vital role in human cardiovascular disease (**Figure 1A and 1B**). Our results also reveal that loss of endothelial CD45 results in a more favorable outcome in an established mouse model of atherosclerosis. In the absence of endothelial CD45, ApoE-null mice fed a WD demonstrate reductions in atherosclerotic lesion size in the aortic sinus, brachiocephalic artery (BCA), and abdominal aorta. We also observed reductions in lesional area throughout the thoracic and abdominal aorta. Overall, these findings plainly indicate that a lack of CD45 from the vascular endothelium inhibits atherosclerosis progression in various arteries.

Although EndoMT contributes to atherosclerotic pathobiology and is associated with more structurally complex plaques^15^, the detailed molecular mechanisms by which CD45 drives EndoMT and the therapeutic potential of manipulating CD45 expression in atherosclerosis remains unknown. Our single-cell RNA-seq analyses provide an unbiased overview of how the transcriptome changes in individual aortic cells^45^. Through single-cell clustering and cell type annotation, we identified the four vascular cell types (**Figure 2A)**: ECs, VSMCs and two distinct cell populations that we designated as EndoMT1 and EndoMT2. In EC-iCD45KO mice, the EC population increases while VSMCs and the EndoMT1 and EndoMT2 populations decrease, which indicates CD45 plays a role in determining vascular cell identity and function.

Analyses of the two EndoMT sub-populations showed that EC-specific CD45 deletion reduces the number of cells in the EndoMT2 sub-population in atherosclerotic mice but had less impact on the EndoMT1 sub-population. Cell identity gene analyses on these two sub-populations showed their strong association with EndoMT-associated GO pathways such as TGFβ signaling, mesenchymal development, and inflammatory signals (**Figure 2K)**. Moreover, the RNA velocity as well as trajectory and cell-cell communication analyses revealed that the EndoMT process is prevalent in WT/ApoE^−/−^ (**Figure 3C to 3D**) and driven by CD45 and TGFβ1 genes (**Figure 3F**). Deletion of CD45 in ECs appears to modulate EndoMT processes by promoting a transition toward MendoT; although, the veracity of this evidence will need to be investigated in future studies using single-cell lineage tracing techniques in mouse models, such as DARLIN and CARLIN^46^.

As noted above, CD45 is not normally expressed in ECs and the details of how it might regulate normal or partial EndoMT remains an open question. This process might be triggered by cross-talk signaling regulatory mechanisms that limit progression through the EndoMT. ECs may undergo progressive transition to a mesenchymal phenotype through intermediate cell states (*i.e.*, partial EndoMT) that enable temporary and reversible adoption of a hybrid endothelial-mesenchymal cell state. It is known that during the early stages of neointima formation, ECs transiently express CD45 accompanied by a host of EndoMT markers^47^. Yamashiro and colleagues also showed that CD31 and α-SMA were expressed in the developing neointima in a carotid artery ligation model^47^.

In this study, human aortic ECs were treated with cobalt chloride (CoCl_­_), a hypoxia-inducible factor (HIF) activator, which resulted in upregulation of CD45 expression^47^. Inhibition of CD45 PTPase activity caused a similar elongated morphology to CoCl_­_treatment; however, the cells treated with CoCl_­_and CD45 PTPase inhibitor showed a significant decrease in VE-Cadherin expression and prominent actin stress fibers. These results suggest that HIF-induced CD45 expression is normally required for the retention of EC fate and cell–cell junctions, and that CD45-positive partial EndoMT contributes to neointima formation and vascular remodeling^47^. While these findings are certainly novel, it is unclear if CD45-positive partial EndoMT is beneficial and more work needs to be done before a mechanism is established.

Nevertheless, our single-cell results comport with these observations as they identified a unique CD45-dependent EndoMT population in the aortic endothelium (EndoMT2 cells) that is seemingly involved in the progressive transition to intermediate mesenchymal cell states in ApoE^−/−^ mice when compared to EC-iCD45KO/ApoE^−/−^ mice fed a WD for 16 weeks. In addition, we observed a reduction of TGFβR2 and TGFβ1 expression using single-cell analysis of the EndoMT2 sub-population from the endothelial CD45-deficient atherosclerotic mice opposed to WT/ApoE^−/−^ mice on a WD for 16 weeks. These observations indicate that endothelial CD45 also mediates partial EndoMT during advanced atherosclerosis^44^.

Previous studies showed that FGFR2 mutations can affect cell responses to TGFβ stimulation via the ERK pathway, which is, at least in part, because of an altered balance between FGF and TGFβ signaling^48^. Consequently, maintaining FGFR2 expression can antagonize TGFβ to restrict EndoMT progression within atherosclerotic lesions. Our study demonstrates that CD45 expression negatively correlates with FGFR2 mRNA expression in MAECs to affect FGFR2 signaling, implying that CD45 inhibition can enhance FGFR2 signaling^48^. As FGFR signaling can obstruct TGFβR-dependent signaling, this may attenuate EndoMT during atherosclerosis.

In line with this hypothesis, it has been reported that FGFR2 overexpression in ECs increases viability, suppresses apoptosis and oxidative stress, and improves endothelial function^49^. In the previously mentioned study on neointima formation^47^, CD45 was detected in cells undergoing EndoMT, and activation of TGFβ signaling was evident in the early neointimal cells as was the expression of Snail, Slug and pSmad2/3^47^. This corresponds with our data where we observed less TGFβR2 and TGFβR3 expression in CD45-deficient MAECs when compared with WT cells. Our findings also show a reduction in the expression of Slug, Twist, MYH9 and Col1a1 in EC-iCD45KO ECs opposed to WT MAECs treated with oxLDL^47^. These findings provide strong evidence that CD45-regulated EndoMT is indispensable for FGFR2 and TGF*β* signal transduction (**Figure 6)**. In addition, preserving KLF2 expression suppresses EndoMT and mitigates endothelial dysfunction during atherosclerosis^50^, which is also consistent with our findings.

In summary, we report the first evidence that in a mouse model of atherosclerosis, the absence of endothelial CD45 leads to reduced lesion development, foam cell infiltration and expression of adhesion molecules in atherosclerosis. Single-cell experiments identified a unique, CD45-dependent EndoMT population in the aortic endothelium (EndoMT2 cells) that mediates partial EndoMT in atherosclerosis^44^. In addition, endothelial CD45 deficiency led to reduced expression of α-SMA and Col1a1, while increasing FGFR2 expression that results in reduced EndoMT and atherosclerotic lesion progression. These findings highlight the potential for therapeutically-targeting endothelial CD45 for the treatment of atherosclerosis, which is the leading cause of death in the United States.

## Author Contributions

Q.P. and H.C. conceived the project. K.A. performed all single-cell data analysis and wrote the original draft of the single-cell data analysis. Q.P. wrote the original draft, performed most of the experiments. Y.W.L. assisted with most of the study. Q.P. and Y.W.L. participated in the experiment design for IF staining and data analysis. B.Z. performed single-cell cDNA library construction. K.C. assisted with *en face* Oil Red O staining and performed a lipid panel test. B.S. assisted with parts of the study, including the studies of MAECs. M.V.M. performed most of the blood counts and lipid panel tests. Q.P. and J.W-S. performed most of the genotype work. J.W-S. assisted with all the genotype work and flow cytometry analysis. S. W. performed the remaining genotype work. Q.P., K.A., J. W-S., K.C., J.B. and H.C. contributed to in-depth discussions of the results. Q.P., K.A., Y.W.L., B.Z., B.S., K.C., J.W-S., S.W., D.B.C., M.A., D-Z.W., K.F.C., J.B., and H.C. reviewed and edited the manuscript.

## Sources of Funding

This project was supported by NIH R01HL146134 (H.C. and D.W.), an American Heart Association Transformational Project Award (H.C.), an American Heart Association Postdoctoral fellowship 23POST1021226 (P.Q.), and, in part, by NIH R01HL156362 (H.C.), R01HL158097 (H.C.), R01HL162367 (H.C.), R01HL174928 (H.C., D.W., and K.F.C.) K99HL171947 (K.F.C.), R01GM125632 (K.F.C.), and R01GM138407 (K.F.C.).

## Data Availability

The authors declare that all supporting data are available within the article and its Supplemental Material. Additional methods or data related to this study are available from the corresponding authors upon reasonable request. The scRNA-seq data generated in this study were deposited into the GEO database with the accession number GSE313399.

## Notes

### Competing Interest Statement

The authors have declared no competing interest.

### Summary of Updates

Revised text, reorganized figures, added new data and authors.

