## Supplemental Materials and Methods for "Novel Role of Endothelial CD45 in Regulating Endothelial-to-Mesenchymal Transition in Atherosclerosis"

#### Genetic Mouse Model

We crossed CD45 floxed mice with tamoxifen-inducible EC-specific Cre deleter mice (iCDH5 CreER$^{T2}$) to create an inducible EC-specific CD45 deficient mouse strain (EC-iCD45KO). We then bred EC-iCD45KO mice to the ApoE$^{-/-}$ (C57BL/6) background to obtain ApoE$^{-/-}$/ECiCD45KO mice. For controls, we used wild-type C57BL/6 mice bearing iCDH5 CreERT2 (denoted as WT). These mice crossed to the ApoE$^{-/-}$ (C57BL/6) background to obtain control ApoE$^{-/-}$/WT (ApoE$^{-/-}$) mice. Mice were fed a Western diet (WD, D12079B, Research Diet) starting at the age of around 8 months old for 8-24 weeks. Mice were sacrificed at different time points based on the experiment.

#### MAEC Isolation

To isolate mouse aortic endothelial cells (MAECs), aortas were collected and washed twice with PBS at 4 °C and then carefully stripped of fat and connective tissue. Aortas were cut into 3 mm long sections, and segments were put on Matrigel-coated (Corning) plate with EC medium. After 4 days, vascular networks were visible under the light microscope and tissue segments were removed. ECs were detached, spun down and cultured in fresh EC medium. The identity of isolated ECs was confirmed by immunofluorescent staining using EC markers CD31. A full list of reagents including antibodies and primers is listed below. Cultured MAECs were treated with 5 μM tamoxifen for 3 days to induce the deletion of CD45 gene from EC-iCD45KO/ApoE$^{-/-}$ mice, MAECs isolated from ApoE$^{-/-}$ mice as controls (without tamoxifen treatment). Cells were treated with 100 μg/mL oxLDL or 10ng/mL TGFβ1 at different times as indicated while maintaining 2 μM tamoxifen in the culture medium.

#### Flow Cytometry

We applied the triple-label flow cytometric method we developed for mitral valve analyses to total aortic ECs from age and gender-matched wild-type (WT) and ApoE$^{-/-}$ mice fed a WD for 16-20 weeks. As ApoE$^{-/-}$ mice fed a normal diet also develop lesions, albeit at a slower rate compared to WD-fed ApoE$^{-/-}$ mice, we chose to use WT mice that were fed a normal chow as our true negative control for EndMT occurrence to rule out the potential effect of high cholesterol content in WD-fed mice and lesion development in normal chow-fed ApoE$^{-/-}$ mice. Blood was thoroughly flushed from aortas to enrich for vessel wall-localized cells. Antibodies were rat anti-mouse CD45 (BD Biosciences), anti-VE-cadherin-FITC (BD Biosciences), anti-CD45-CF647 (BD Biosciences), anti-FGFR1-PE (BD Biosciences) and rabbit anti-human VE-cadherin (ABD Serotec) coupled to APC using the LYNX Rapid APC Antibody Conjugation Kit (ABD Serotec) as well as rabbit anti-human α-SMA-PE (Abcam) with isotype-matched control IgGs and matching fluorescent tags. Antibodies were authenticated by showing binding to murine CD45, VE-cadherin, and α-SMA, respectively, and using appropriate positive and negative controls.

### SUPPLEMENTAL TABLE 1: ANTIBODIES

| Target antigen | Vendor or Source | Catalog # | Working concentration |
|---|---|---|---|
| Primary Antibodies | | | |
| Mouse VE-Cadherin Antibody | R & D systems | AF1002 | 1:50 (IF) |
| Anti-Klf2 Antibody (Rabbit Polyclonal Antibody | EMD Millipore | 09-820 | 1:200 (IF) |
| Mouse monoclonal anti-CD68 | Santa Cruz Biotechnology | Sc20060 | 1:200 (IF) |
| Anti-Actin, α-Smooth Muscle - Cy3™ antibody, Mouse monoclonal α-SMA | Sigma-Aldrich | C6198 | 1:200(IF) |
| αSMA | Millipore Sigma | A-2547 | 137.5 ug/ml |
| Rat-anti Mouse CD31 | BD Pharmingen | 550274 | 1:50 (IF) |
| Mouse Anti-CD45 | Santa Cruz Biotechnology | Sc-28369 | 1:200 |
| Mouse monoclonal ICAM-1 | Santa Cruz Biotechnology | Sc8439 | 1:200 (IF) |
| Mouse monoclonal VCAM-1 | Santa Cruz Biotechnology | sc13160 | 1:70 (IF) |

| Alexa Fluor Conjugated Second Antibodies | | | |
| --- | --- | --- | --- |
| Goat anti-rabbit 647 | Invitrogen | A-21244 | 1:200 (IF) |
| Donkey anti-rat 488 | Invitrogen | A-21208 | 1:200 (IF) |
| Donkey anti-goat 594 | Invitrogen | A11058 | 1:200 (IF) |
| Donkey anti-mouse 647 | Invitrogen | A31571 | 1:200 (IF) |

### SUPPLEMENTAL TABLE 2: REAGENTS

| Description | Source / Repository | Catalog # |
|---|---|---|
| Oil Red O (ORO) | Thermo Scientific | A12989 |
| Hematoxylin and Eosin stain kit | Vector Laboratories | H-3502 |
| Propylene glycol | VWR | 0575 |
| Oxidized LDL (oxLDL) | Athens Research and Technology | 12-16-120412-ox |
| Low Density Lipoprotein from Human Plasma, oxidized, DiI conjugate (DiI- OxLDL) | ThermoFisher Scientific | L34358 |
| SlowFade mount with DAPI | Invitrogen | 1896320 |
| Protein inhibitor cocktail | Complete | REF:11836170001 |
| Isopropanol, molecular biology grade | ThermoFisher Scientific | T036181000CS |
| Qiagen RNeasy Mini Kit | Qiagen | 74104 |
| RNase-free DNAse Set | Qiagen | 79254 |
| HiScript II One Step qRT-PCR SYBR Green | Vazyme | Q221-01 |
| SYPR Green qPCR Master Mix reagent | Vazyme | Q712 |
| Fluoroshield | R & D Systems | F6812 |

| | | |
|---|---|---|
| Ciprofloxacin Hydrochloride | TCI AMERICA | C2227 |
| Triton™ X-100 Surfactant | MilliporeSigma | TX15681 |
| Matrigel Matrix | Corning | 356237 |
| CD31 MicroBeads (Mouse) | Miltenyi Biotec | 130-097-418 |
| Collagenase type I | Gibco by Life technologies | 17100-017 |
| Collagenase type IV | Gibco by Life technologies | 17104-019 |
| Liberase | Roche | 05401127001 |
| GEM Single Cell 3’GEM Kit v3.1 | 10X Genomics | PN-1000123 |
| GEM Single Cell 3’Library Kit v3.1 | 10X Genomics | PN-1000157 |
| GEM Single Cell 3’Gel Bead Kit v3.1 | 10X Genomics | PN-1000122 |
| Dynabeads™ MyOne™ SILANE | 10X Genomics | PN-2000048 |
| GEM Chip G Single Cell Kit, 48rxns | 10X Genomics | PN-1000120 |
| Single Index Kit T Set A, 96 rxns | 10X Genomics | PN-1000213 |

### SUPPLEMENTAL TABLE 3: CELL CULTURE SUPPLIES

| Name | Vendor or Source | Catalog # |
|---|---|---|
| DMEM | Corning | 10-013-CV |
| F12K | Corning | 10-025-CV |
| FBS | Phoenix Scientific | PS-300 |
| Ciprofloxacin Hydrochloride | TCI America | C2227 |
| Recombinant Mouse TGF-beta 1 Protein | R&D Systems | 7346-B2/C |
| Heparin | Sigma | H4784-1G |
| Pyruvate | Gibco | 11360-070 |
| Gibco Penicillin Streptomycin | ThermoFisher | 10378016 |

### SUPPLEMENTAL TABLE 4: OLIGONUCLEOTIDES

| Description | Source / Repository |
|---|---|
| TGFβR2 | This study |
| Forward: 5'- GGACCCTACTCTGTCTGTGG -3'<br>Reverse: 5'- AGCCATGGAGTAGACATCCG -3' | |
| TGFβR3 | This study |
| Forward: 5'- GACATCCCTTCCACCCAAGA-3'<br>Reverse: 5'- CAGGAGGAATGGTGTGGACT -3' | |
| Beta-actin | This study |
| Forward: 5'- GATCAAGATCATTGCTCCTCCTG -3'<br>Reverse: 5'- AGGGTGTAAAACGCAGCTCA-3' | |
| FGFR2 | This study |
| Forward:5'TTTAGAACCAGAAGAGCCACCA-3'<br>Reverse: 5' - ATCACGGCGGCATCTTTCA -3' | |
| Slug | This study |
| Forward: 5-ACTACAGCGAACTGGACACA<br>Reverse: 5-GCCACTGGGTAAAGGAGAGT | |

| | |
|---|---|
| Twist1 | This study |
| Forward: 5-GACTCCAAGATGGCAAGCTG<br>Reverse :5-GCCACTGGGTAAAGGAGAGT | |
| vWF | This study |
| Forward: 5-GCCAGTATGTTCTGGTGCAG<br>Reverse :5-CGTTCACCTCTCCGTCAAAC | |
| Collagen 1a (Col1a1)3 | This study |
| Forward: 5-AGGGTGACAAAGGAGAGGTG<br>Reverse: 5-AGGTGGAAGAGGTCAATGGG | |
| VE-Cadherin | This study |
| Forward: 5- TACCACTTCAAGCTGCCAGA<br>Reverse: 5-TCGGAAGAATTGGCCTCTGT | |

### SUPPLEMENTAL TABLE 5: SOFTWARE AND ALGORITHMS

| Description | Persistent ID / URL |
|---|---|
| Image J | https://imagej.nih.gov/ij/ |
| GraphPad Prism | https://www.graphpad.com/scientificsoftware/prism/ |
| Zeiss Zen lite | https://www.zeiss.com/microscopy/us/products/microscope software/zen-lite.html |
| Seuart | https://github.com/satijalab/seurat |
| MONOCLE | https://github.com/cole-trapnell-lab/monocle3 |
| ScVelo | https://github.com/theislab/scvelo/tree/main |
| CellRank | https://github.com/theislab/cellrank |
| ClusterProfiler | https://github.com/YuLab-SMU/clusterProfiler |
| NicheNet | https://github.com/saeyslab/nichenetr |
| SCIG | https://zenodo.org/records/14726426 |

#### Data Availability

| Description | Source / Repository | Persistent ID / URL |
|---|---|---|
| ScRNA seq mice models | GEO | GSE313399 |
