## Supplemental Figures for "Novel Role of Endothelial CD45 in Regulating Endothelial-to-Mesenchymal Transition in Atherosclerosis"

#### Figure S1

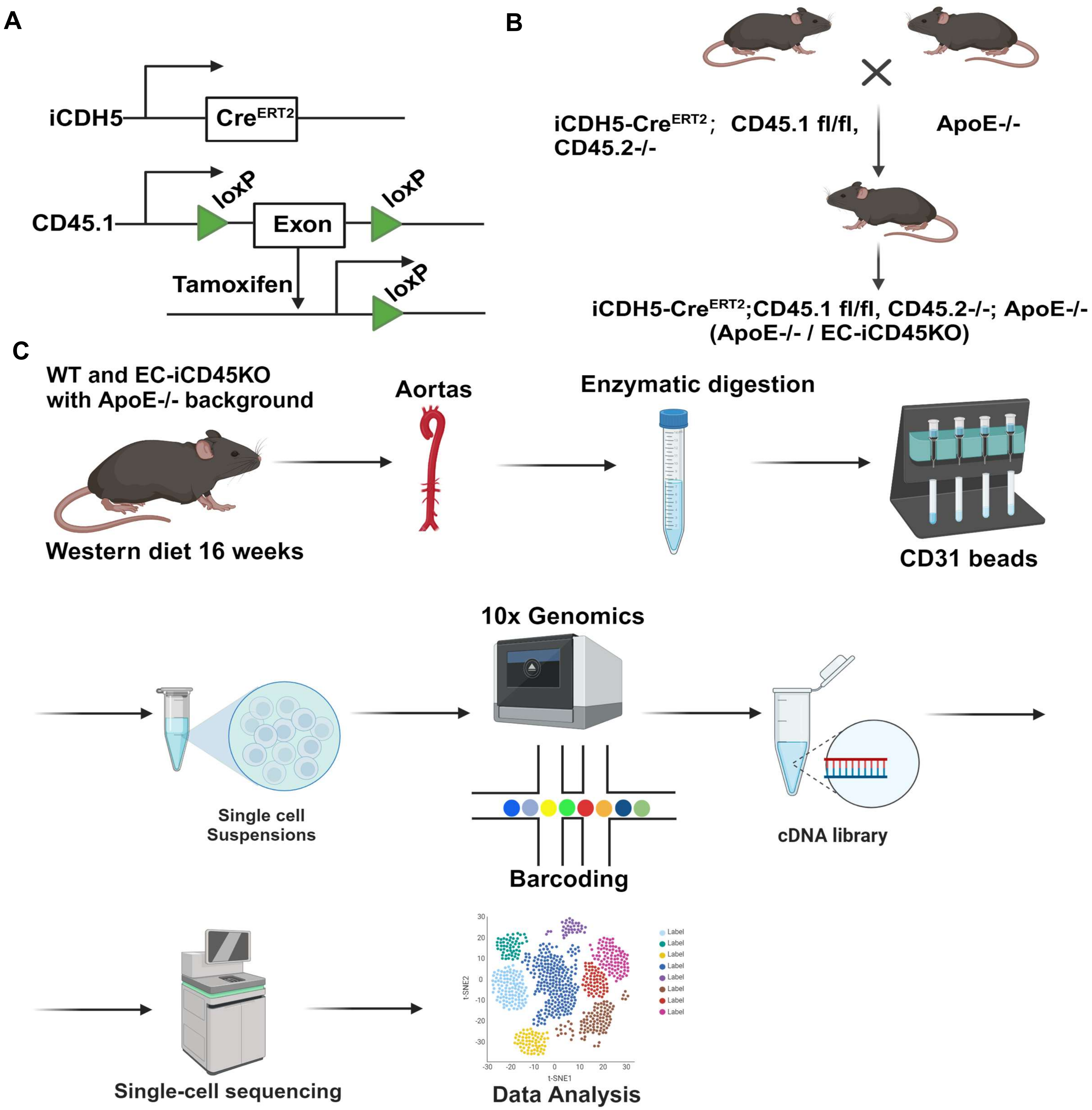
A
iCDH5
CreERT2
CD45.1
loxP
Exon
loxP
Tamoxifen
loxP
B
iCDH5-CreERT2; CD45.1 fl/fl, CD45.2-/-
ApoE-/-
iCDH5-CreERT2;CD45.1 fl/fl, CD45.2-/-; ApoE-/-
(ApoE-/- / EC-iCD45KO)
C
WT and EC-iCD45KO with ApoE-/- background
Western diet 16 weeks
Aortas
Enzymatic digestion
CD31 beads
Single cell Suspensions
10x Genomics
Barcoding
cDNA library
Single-cell sequencing
Data Analysis
t-SNE1
t-SNE2

**Figure S2**

| Parameters | WT/ApoE$^{-/-}$ mice (n=11) | EC-iCD45KO/ApoE$^{-/-}$ mice (n=9) | P value |
|---|---|---|---|
| Cholesterol(mg/dL) | 800.7± 118.2 | 834.7 ± 113.3 | 0.6700 |
| Triglycerides(mg/dL) | 124.2.8±28.46 | 103.6 ± 19.02 | 0.8820 |
| LDL (mg/dL) | 728.6 ± 111.9 | 762.4 ± 117.4 | 0.5516 |
| HDL(mg/dL) | 73.27 ± 13.99 | 79.44 ± 18.03 | 0.9270 |

### Figure S3

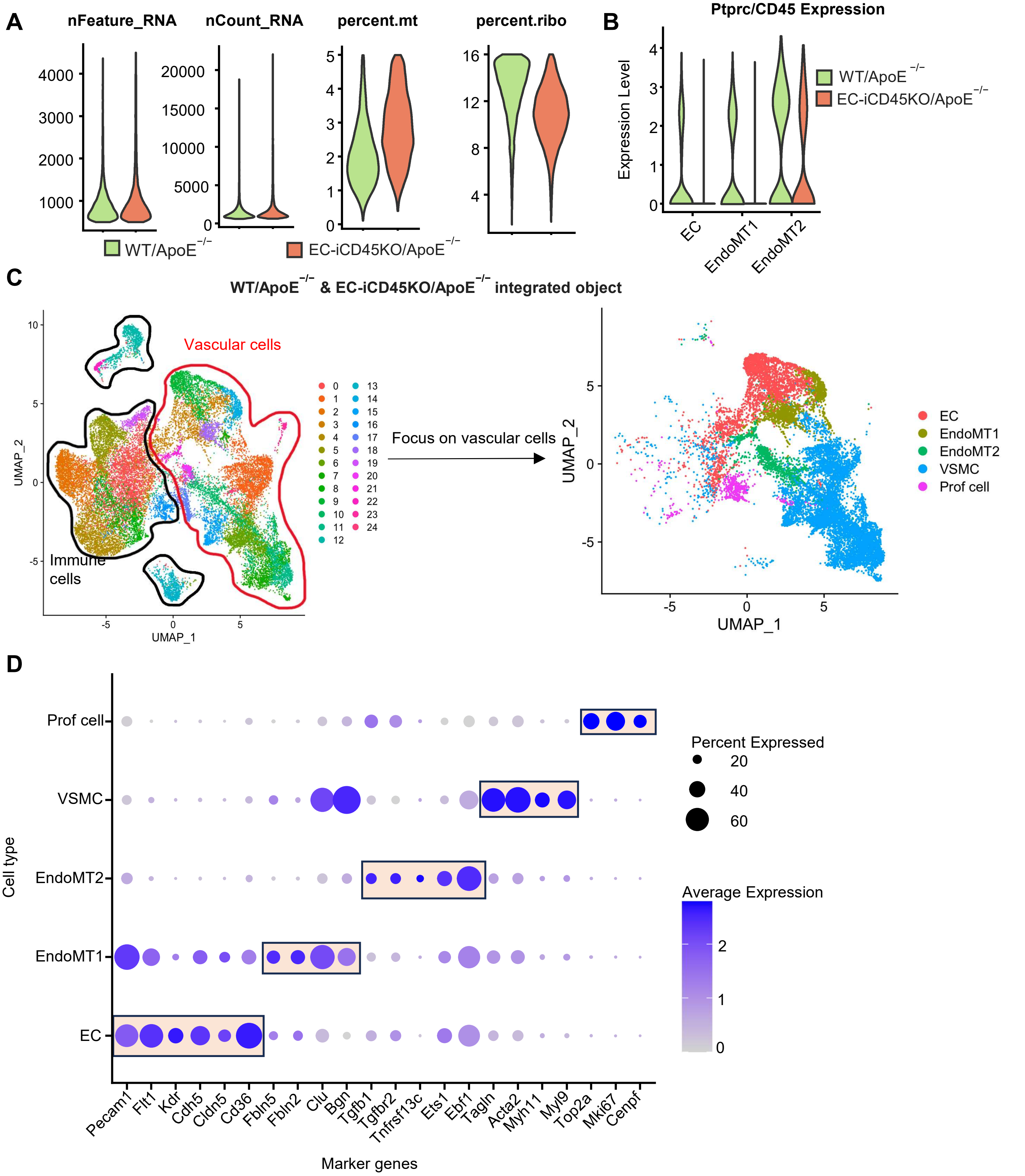

A
nFeature_RNA
nCount_RNA
percent.mt
percent.ribo
WT/ApoE−/−
EC-iCD45KO/ApoE−/−
B
Ptprc/CD45 Expression
Expression Level
EC
EndoMT1
EndoMT2
C
WT/ApoE−/− & EC-iCD45KO/ApoE−/− integrated object
Vascular cells
Immune cells
Focus on vascular cells
UMAP_1
UMAP_2
EC
EndoMT1
EndoMT2
VSMC
Prof cell
D
Prof cell
VSMC
EndoMT2
EndoMT1
EC
Cell type
Percent Expressed
20
40
60
Average Expression
Pecam1
Flt1
Kdr
Cdh5
Cldn5
Cd36
Fbln5
Fbln2
Clu
Bgn
Tgfb1
Tgfbr2
Tnfrsf13c
Ets1
Ebf1
Tagln
Acta2
Myh11
Myl9
Top2a
Mki67
Cenpf
Marker genes

### Figure S4

A

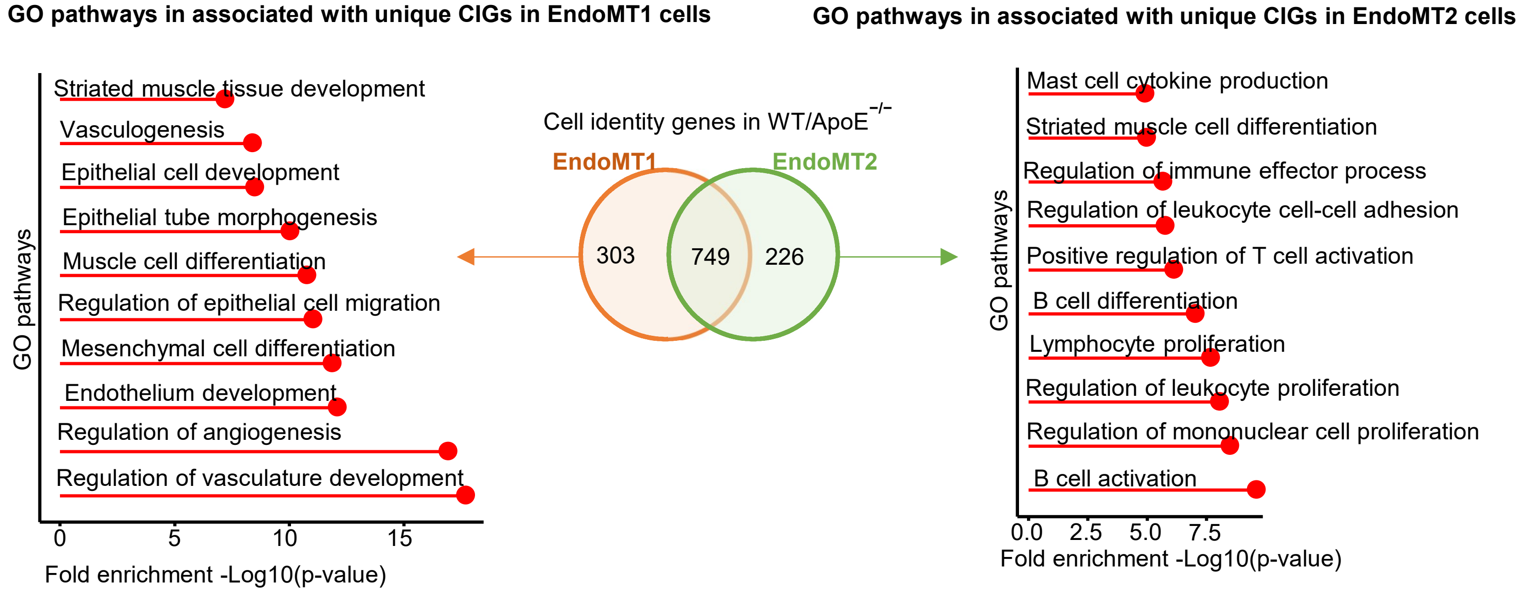
GO pathways in associated with unique CIGs in EndoMT1 cells
GO pathways in associated with unique CIGs in EndoMT2 cells
Striated muscle tissue development
Vasculogenesis
Epithelial cell development
Epithelial tube morphogenesis
Muscle cell differentiation
Regulation of epithelial cell migration
Mesenchymal cell differentiation
Endothelium development
Regulation of angiogenesis
Regulation of vasculature development
GO pathways
0
5
10
15
Fold enrichment -Log10(p-value)
Cell identity genes in WT/ApoE−/−
EndoMT1
EndoMT2
303
749
226
Mast cell cytokine production
Striated muscle cell differentiation
Regulation of immune effector process
Regulation of leukocyte cell-cell adhesion
Positive regulation of T cell activation
B cell differentiation
Lymphocyte proliferation
Regulation of leukocyte proliferation
Regulation of mononuclear cell proliferation
B cell activation
0.0
2.5
5.0
7.5
Fold enrichment -Log10(p-value)

B

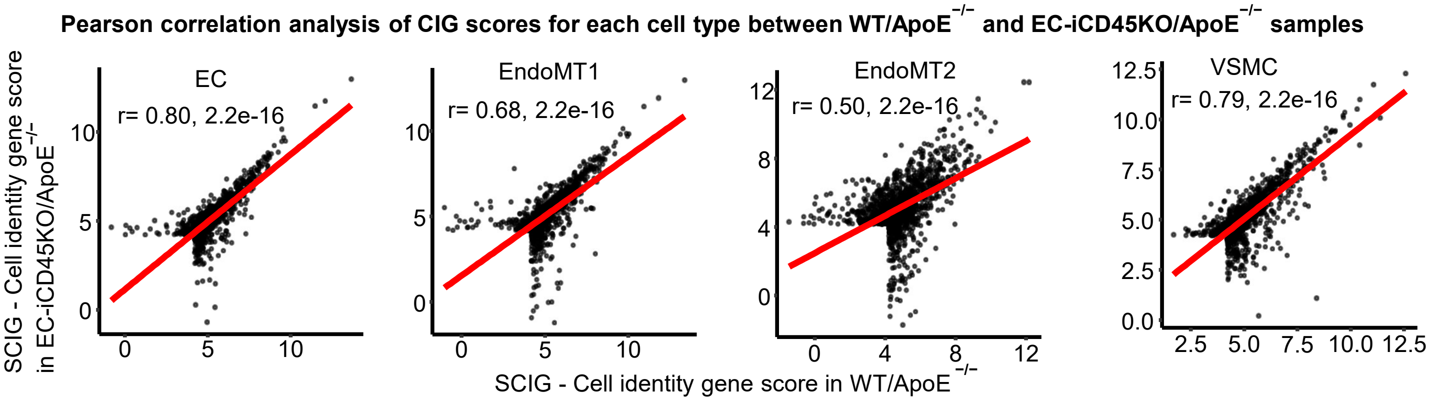
Pearson correlation analysis of CIG scores for each cell type between WT/ApoE−/− and EC-iCD45KO/ApoE−/− samples
SCIG - Cell identity gene score in EC-iCD45KO/ApoE−/−
EC
r= 0.80, 2.2e-16
EndoMT1
r= 0.68, 2.2e-16
EndoMT2
r= 0.50, 2.2e-16
VSMC
r= 0.79, 2.2e-16
SCIG - Cell identity gene score in WT/ApoE−/−

### Figure S5

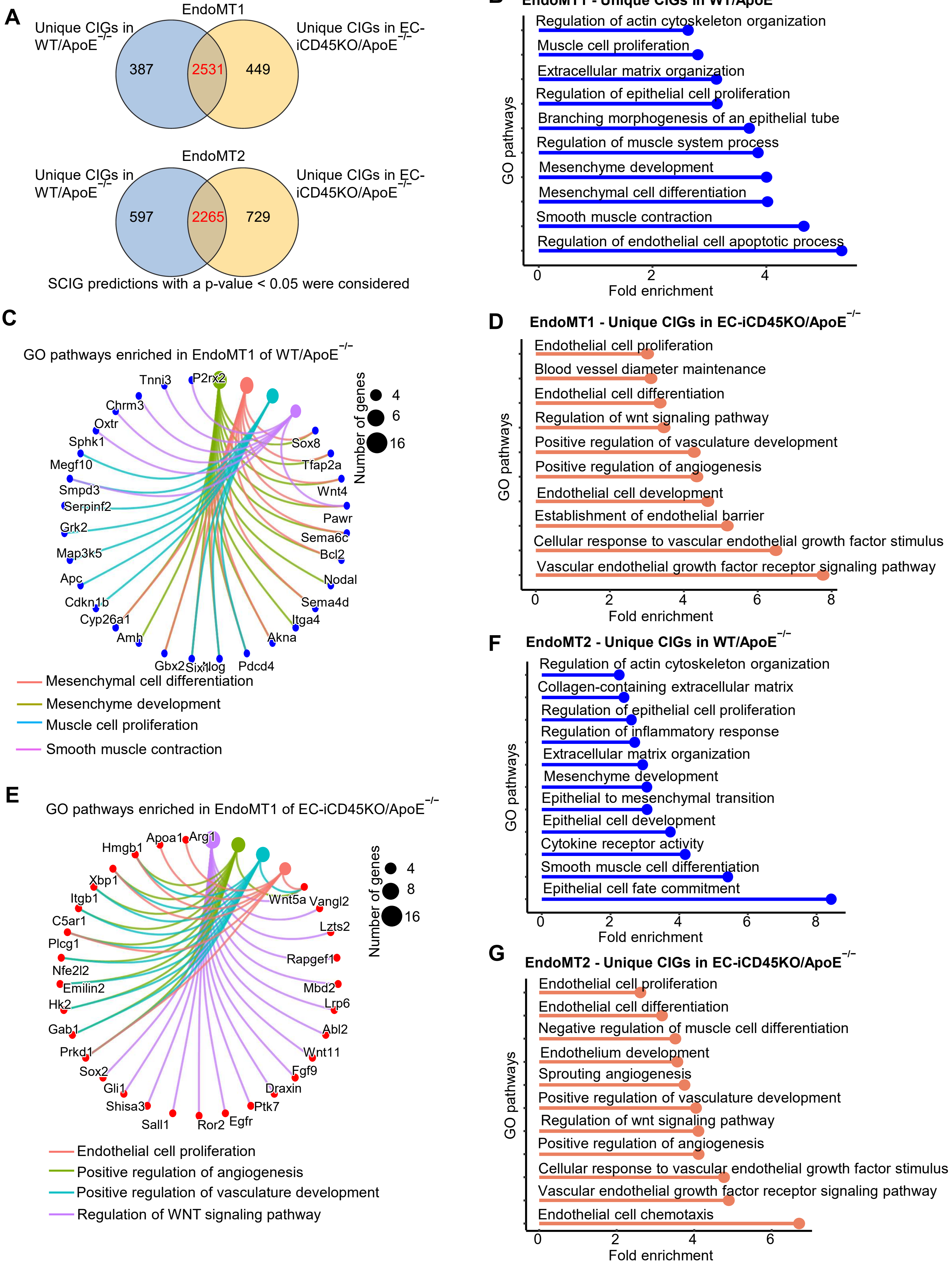

A
EndoMT1
Unique CIGs in WT/ApoE−/−
Unique CIGs in EC-iCD45KO/ApoE−/−
387
2531
449
EndoMT2
Unique CIGs in WT/ApoE−/−
Unique CIGs in EC-iCD45KO/ApoE−/−
597
2265
729
SCIG predictions with a p-value < 0.05 were considered
B
EndoMT1 - Unique CIGs in WT/ApoE−/−
Regulation of actin cytoskeleton organization
Muscle cell proliferation
Extracellular matrix organization
Regulation of epithelial cell proliferation
Branching morphogenesis of an epithelial tube
Regulation of muscle system process
Mesenchyme development
Mesenchymal cell differentiation
Smooth muscle contraction
Regulation of endothelial cell apoptotic process
GO pathways
Fold enrichment
0
2
4
C
GO pathways enriched in EndoMT1 of WT/ApoE−/−
Tnni3
P2rx2
Chrm3
Oxtr
Sphk1
Megf10
Smpd3
Serpinf2
Grk2
Map3k5
Apc
Cdkn1b
Cyp26a1
Amh
Gbx2
Six1
Hog
Pdcd4
Akna
Itga4
Sema4d
Nodal
Bcl2
Sema6c
Pawr
Wnt4
Tfap2a
Sox8
Number of genes
4
6
16
Mesenchymal cell differentiation
Mesenchyme development
Muscle cell proliferation
Smooth muscle contraction
D
EndoMT1 - Unique CIGs in EC-iCD45KO/ApoE−/−
Endothelial cell proliferation
Blood vessel diameter maintenance
Endothelial cell differentiation
Regulation of wnt signaling pathway
Positive regulation of vasculature development
Positive regulation of angiogenesis
Endothelial cell development
Establishment of endothelial barrier
Cellular response to vascular endothelial growth factor stimulus
Vascular endothelial growth factor receptor signaling pathway
GO pathways
Fold enrichment
0
2
4
6
8
E
GO pathways enriched in EndoMT1 of EC-iCD45KO/ApoE−/−
Hmgb1
Apoa1
Arg1
Xbp1
Itgb1
C5ar1
Plcg1
Nfe2l2
Emilin2
Hk2
Gab1
Prkd1
Sox2
Gli1
Shisa3
Sall1
Ror2
Egfr
Ptk7
Draxin
Fgf9
Wnt11
Abl2
Lrp6
Mbd2
Rapgef1
Lzts2
Vangl2
Wnt5a
Number of genes
4
8
16
Endothelial cell proliferation
Positive regulation of angiogenesis
Positive regulation of vasculature development
Regulation of WNT signaling pathway
F
EndoMT2 - Unique CIGs in WT/ApoE−/−
Regulation of actin cytoskeleton organization
Collagen-containing extracellular matrix
Regulation of epithelial cell proliferation
Regulation of inflammatory response
Extracellular matrix organization
Mesenchyme development
Epithelial to mesenchymal transition
Epithelial cell development
Cytokine receptor activity
Smooth muscle cell differentiation
Epithelial cell fate commitment
GO pathways
Fold enrichment
0
2
4
6
8
G
EndoMT2 - Unique CIGs in EC-iCD45KO/ApoE−/−
Endothelial cell proliferation
Endothelial cell differentiation
Negative regulation of muscle cell differentiation
Endothelium development
Sprouting angiogenesis
Positive regulation of vasculature development
Regulation of wnt signaling pathway
Positive regulation of angiogenesis
Cellular response to vascular endothelial growth factor stimulus
Vascular endothelial growth factor receptor signaling pathway
Endothelial cell chemotaxis
GO pathways
Fold enrichment
0
2
4
6

### Figure S6

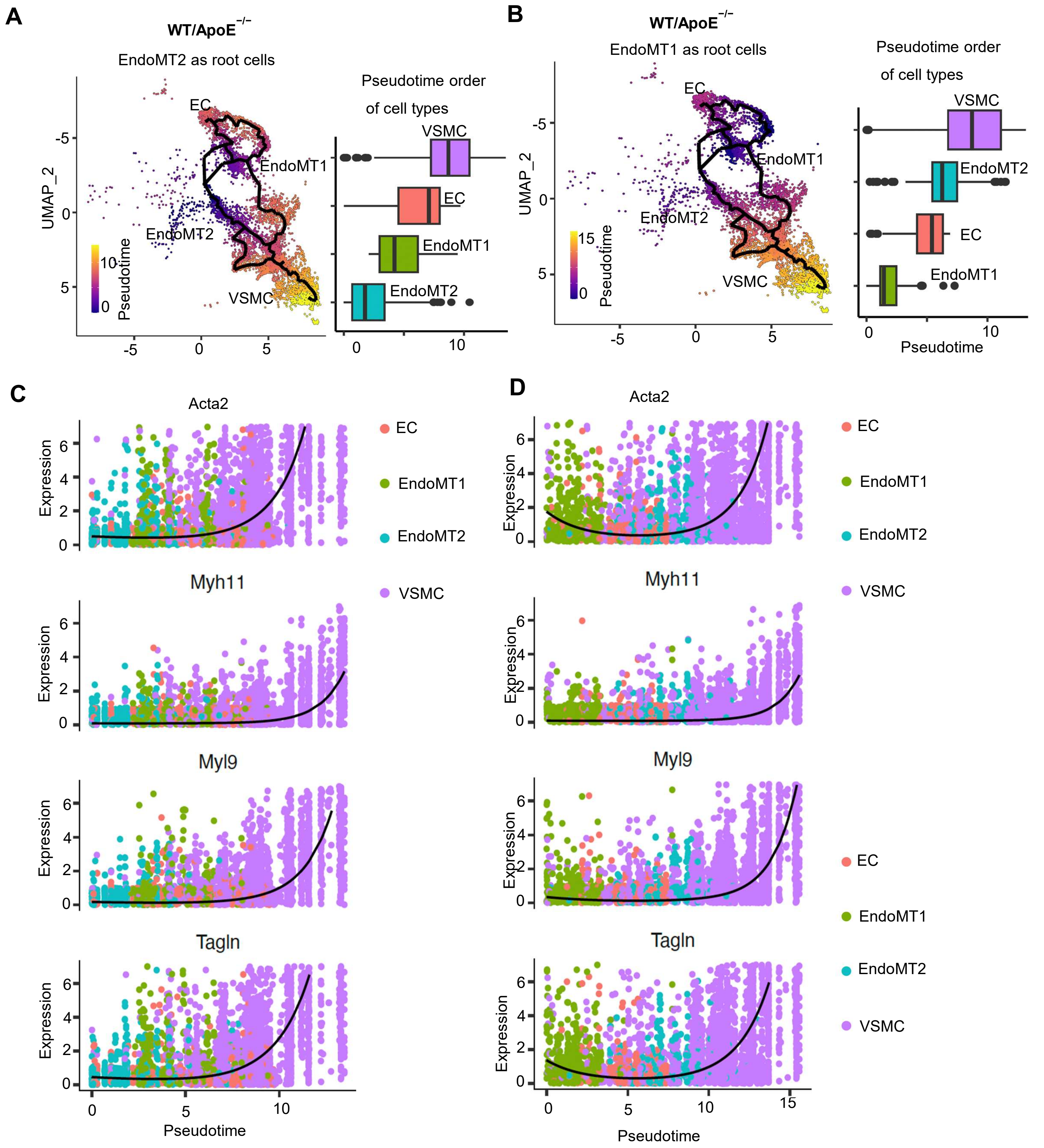
A
WT/ApoE−/−
EndoMT2 as root cells
UMAP_2
Pseudotime
EC
EndoMT1
EndoMT2
VSMC
Pseudotime order
of cell types
B
WT/ApoE−/−
EndoMT1 as root cells
C
Acta2
Myh11
Myl9
Tagln
Expression
Pseudotime
D

### Figure S7

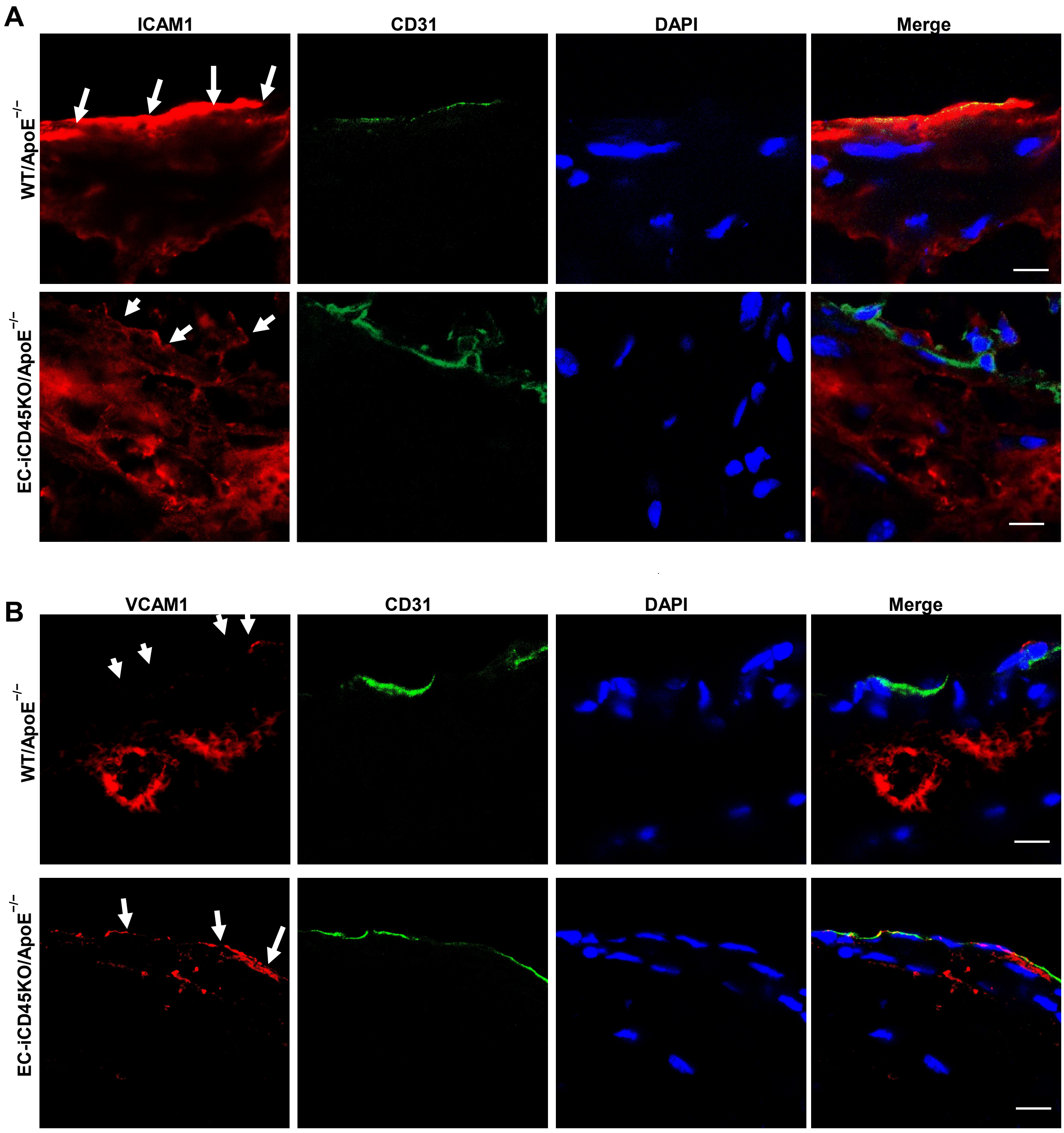
A
ICAM1
CD31
DAPI
Merge
WT/ApoE$^{-/-}$
EC-iCD45KO/ApoE$^{-/-}$
B
VCAM1
CD31
DAPI
Merge
WT/ApoE$^{-/-}$
EC-iCD45KO/ApoE$^{-/-}$

### Figure S8

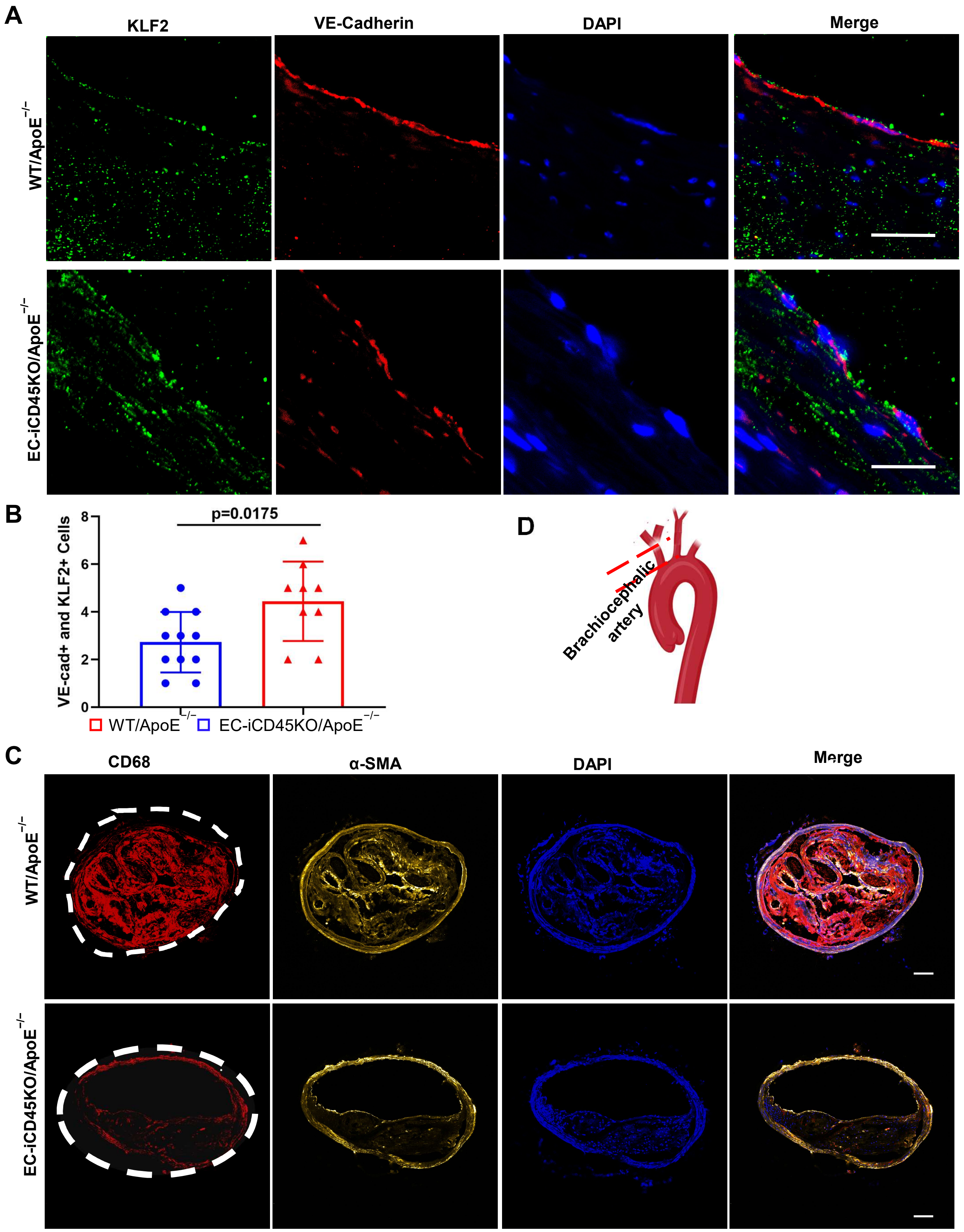
A
KLF2
VE-Cadherin
DAPI
Merge
WT/ApoE−/−
EC-iCD45KO/ApoE−/−
B
p=0.0175
VE-cad+ and KLF2+ Cells
0
2
4
6
8
WT/ApoE−/−
EC-iCD45KO/ApoE−/−
D
Brachiocephalic artery
C
CD68
α-SMA
DAPI
Merge
WT/ApoE−/−
EC-iCD45KO/ApoE−/−

### Figure S9

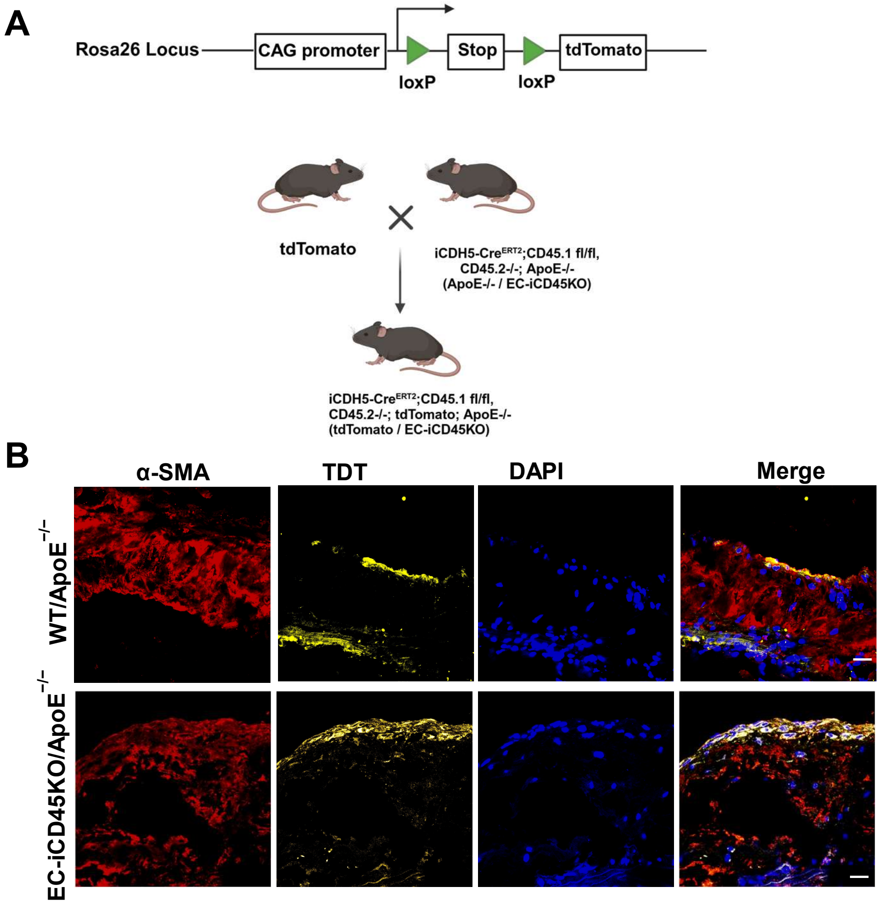
A
Rosa26 Locus
CAG promoter
Stop
tdTomato
loxP
loxP
tdTomato
iCDH5-CreERT2;CD45.1 fl/fl,
CD45.2-/-; ApoE-/-
(ApoE-/- / EC-iCD45KO)
iCDH5-CreERT2;CD45.1 fl/fl,
CD45.2-/-; tdTomato; ApoE-/-
(tdTomato / EC-iCD45KO)
B
α-SMA
TDT
DAPI
Merge
WT/ApoE−/−
EC-iCD45KO/ApoE−/−

### Figure S10

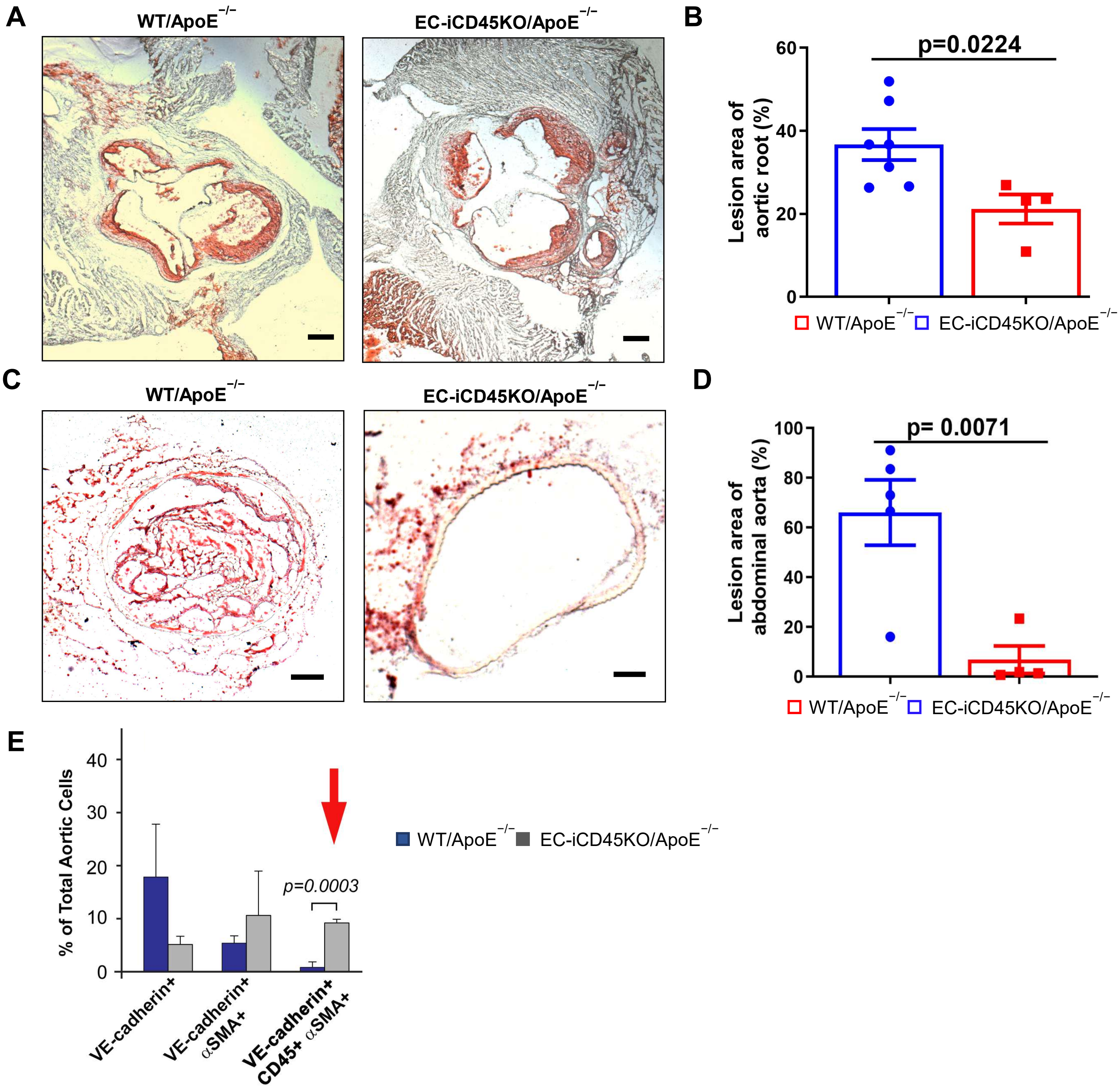

A
WT/ApoE−/−
EC-iCD45KO/ApoE−/−
B
p=0.0224
Lesion area of aortic root (%)
60
40
20
0
WT/ApoE−/−
EC-iCD45KO/ApoE−/−
C
WT/ApoE−/−
EC-iCD45KO/ApoE−/−
D
p= 0.0071
Lesion area of abdominal aorta (%)
100
80
60
40
20
0
WT/ApoE−/−
EC-iCD45KO/ApoE−/−
E
% of Total Aortic Cells
40
30
20
10
0
p=0.0003
WT/ApoE−/−
EC-iCD45KO/ApoE−/−
VE-cadherin+
VE-cadherin+ αSMA+
VE-cadherin+ CD45+ αSMA+

### Figure S11

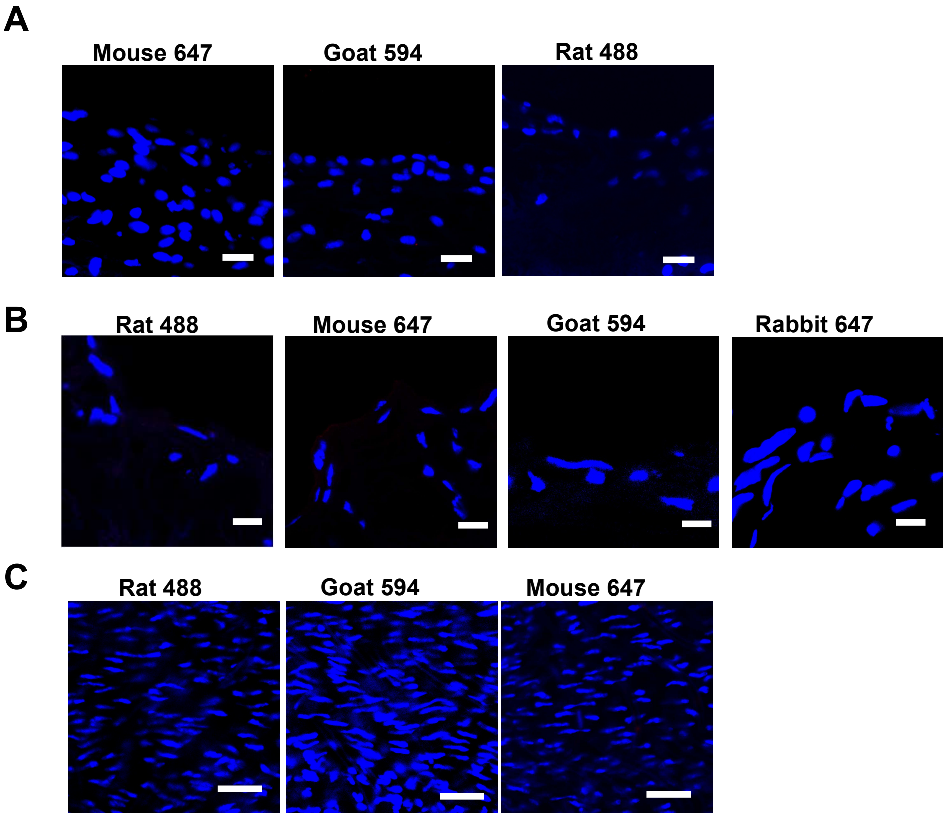
A
Mouse 647
Goat 594
Rat 488
B
Rat 488
Mouse 647
Goat 594
Rabbit 647
C
Rat 488
Goat 594
Mouse 647
