## Supplemental Figure Legends for "Novel Role of Endothelial CD45 in Regulating Endothelial-to-Mesenchymal Transition in Atherosclerosis"

**Figure S1. Overview of the animal model and pipeline for 10X genomics scRNA-seq. A**,**B**, Workflow for generation of CD45.1$^{fl/fl}$ CD45.2$^{-/-}$ (EC-iCD45KO)/ApoE$^{-/-}$ mice with WT/ApoE$^{-/-}$ mice as the control group. **C**, Workflow illustrating the steps involved in single-cell RNA sequencing (scRNA-seq), including sample preparation, sequencing, and data analysis. WT and EC-iCD45KO male mice were fed a Western diet (WD) for 16 weeks, followed by the isolation of aortas, enzyme digestion, CD31 bead enrichment of endothelial cells (ECs). Single cell suspensions were prepared according to the manufacturer's instructions (10x Genomics). Single-cell cDNA libraries were prepared as follows: 1) barcoding cells for 10x Genomics, 2) cDNA library construction and 3) scRNA sequencing. cDNA library sequencing and computational analysis were performed by preprocessing sequencing reads by Cell Ranger (10x Genomics) prior to downstream analysis.

**Figure S2. Serum lipid profile for WT/ApoE$^{-/-}$ and EC-iCD45KO/ApoE$^{-/-}$ mice.** The levels of cholesterol, triglycerides, LDL and HDL are shown (mg/dL). There was no significant difference in P values between WT/ApoE$^{-/-}$ (n = 11) and EC-iCD45KO/ApoE$^{-/-}$ (n = 9) mice.

**Figure S3. Data processing, quality control, and cell clustering analysis of scRNA-seq from WT/ApoE$^{-/-}$ and EC-iCD45KO/ApoE$^{-/-}$ mice. A**, High-quality cells were selected from 11,497 WT/ApoE$^{-/-}$ and 16,273 EC-iCD45KO/ApoE$^{-/-}$ CD31-enriched cells based on nFeatureRNA, nCountRNA, and mitochondrial/ribosomal gene percentages, resulting in 6,476 and 7,021 cells, respectively. **B**, Violin plot showing the expression levels of CD45/PTPRC across the identified cell types. **C**, Extraction of cell clusters expressing EC, VSMC, EndoMT, and proliferative cell markers in WT/ApoE$^{-/-}$ and EC-iCD45KO/ApoE$^{-/-}$ mice. **D**, Dot plot visualization of marker gene expression in the five identified cell types.

**Figure S4: Characterization of cell identity genes in EndoMT1 and EndoMT2 cells in WT/ApoE$^{-/-}$ and CD45-deletion mice. A**, Unique cell identity genes of EndoMT1 and EndoMT2 cells identified using SCIG, along with their Gene Ontology (GO) pathway enrichment analysis. **B**, Pearson correlation of cell identity gene (CIG) scores for each cell type in WT/ApoE$^{-/-}$ and EC-iCD45KO/ApoE$^{-/-}$ mice.

**Figure S5. Cell-identity gene and functional pathway analyses reveal endothelial CD45 deletion reshapes EndoMT1 and EndoMT2 functional programs. A**, Venn diagram comparing cell-identity genes (CIGs) in EndoMT1 and EndoMT2 cells, highlighting shared and unique genes in WT/ApoE$^{-/-}$ and EC-iCD45KO/ApoE$^{-/-}$ mice. **B**,**C**, Gene Ontology (GO) pathway enrichment analysis of unique EndoMT1 CIGs in WT/ApoE$^{-/-}$ mice shown as a lollipop plot (B) and a gene-concept network diagram illustrating relationships between enriched pathways and associated genes (C). **D**,**E**, GO pathway enrichment analysis of upregulated EndoMT2 CIGs in EC-iCD45KO/ApoE$^{-/-}$ mice, displayed as a lollipop plot (D) and a gene-concept network diagram (E). **F,G**, Lollipop plots showing enriched pathways of unique EndoMT1 CIGs in WT/ApoE$^{-/-}$ (F) and EC-iCD45KO/ApoE$^{-/-}$ (G) mice.

**Figure S6. Monocle trajectory analysis reveals that EndoMT is prevalent in WT/ApoE$^{-/-}$ and EC-iCD45KO/ApoE$^{-/-}$ mice. A**,**B**, Monocle trajectory analysis using EndoMT1 (A) and EndoMT2 (B) cells showing transitions toward VSMCs. **C,D**, Expression of VSMC marker genes aligned with EndoMT trajectory when EndoMT1 (C) or EndoMT2 (D) cells are used as the root cell.

**Figure S7. Loss of endothelial CD45 reduces cell adhesion molecular expression in atherosclerotic mice. A**, ICAM-1 (red), CD31 (green) and DAPI (blue) on the luminal surface of aortic root sections (16-week WD). **B**, VCAM-1 (red), CD31 (green) and DAPI (blue) on the luminal surface of aortic root sections (12-week WD) (Scale bars = 10 μm).

**Figure S8. Loss of endothelial CD45 increases KLF-2 colocalization with VE-Cadherin in the aortic root and decreases αSMA expression in the brachiocephalic artery. A**, KLF-2 (green), VE-Cadherin (red) and DAPI (blue) on the luminal surface of aortic root sections (12-week WD) (Scale bars = 10 μm). **B**, Quantifications for co-stained KLF-2 and VE-Cadherin. **C**, CD68 (red), αSMA (yellow) and DAPI (blue) on the luminal surface of BCA sections (16-week WD) (Scale bars = 100 μm). **D**, The structure of the brachiocephalic artery (BCA).

**Figure S9. Loss of endothelial CD45 inhibits EndoMT *in vitro* and *in vivo*. A**, Workflow of tdTomato, CD45.1$^{fl/fl}$, CD45.2$^{-/-}$/ApoE$^{-/-}$ and tdTomato/EC-iCD45KO/ApoE$^{-/-}$ mice. **B**, Immunofluorescence staining of aortic root sections from tdTomato/ApoE$^{-/-}$ and tdTomato/EC-iCD45KO/ApoE$^{-/-}$ (13-week WD) with anti-αSMA. (S7B, Scale bars = 20 μm).

**Figure S10. Loss of endothelial CD45 reduces atherosclerosis in WT/ApoE$^{-/-}$ mice. A-D**, Aortic root (A), and aortic aneurysm sections (C) from WT/ApoE$^{-/-}$ and EC-iCD45KO mice (16-week WD) stained with Oil Red O (ORO). Quantifications for *en face* ORO staining areas in aortic root (B) and aortic aneurysm sections (D). **E**, The arrow shows CD45$^{+}$/VE-cadherin$^{+}$/αSMA$^{+}$ cells were significantly increased from 0.8 ± 1.0% to 9.2 ± 0.7% in WT/ApoE$^{-/-}$ WD-fed mice (n=3) ($p = 0.0003$). The presence of α-SMA$^{+}$ in WT/ApoE$^{-/-}$ WD-fed mice indicates CD45$^{+}$ ECs were undergoing EndoMT in aortic atherosclerotic lesions.

**Figure S11. Staining with isotype-matched control IgGs, the corresponding fluorescently tagged secondary antibodies and DAPI. A**, Isotype-matched control IgGs for paraffin-embedded sections. **B-D**, Representative staining of cryosections (B) thoracic sections (C), and MAECs (D). Scale bars = 10 μm.
